# PyiTOL: reproducible Python workflows for iTOL annotation and taxonomic monophyly assessment

**DOI:** 10.64898/2026.08.27.747471

**Authors:** Zichao Zeng, Yinzhao Wang

## Abstract

**Motivation:** The Interactive Tree of Life (iTOL) is widely used to display and annotate phylogenetic trees, but managing its format-sensitive annotation files impede reproducible high-throughput analyses. Among the maintained Python packages and versions evaluated, none combined template generation, taxonomic monophyly assessment and iTOL batch operations.

**Results:** PyiTOL validates inputs, generates 31 iTOL template schemas (22 accepted by the live batch uploader), performs LCA-based monophyly classification with nested-monophyly detection, sampling-completeness states and polyphyletic subgroup decomposition, plus API upload and session replay. On a topology-constructed benchmark, all calls matched prespecified labels for 4,389 groups; on a 700-genome tree, binary mono/non-mono calls agreed with ETE4 for 409 genera; 17,294 GTDB R232 genera were processed in about 17 s.

**Availability and Implementation:** PyiTOL 1.0.3 (Python ≥3.10; Linux, macOS and Windows) is MIT-licensed at https://github.com/ZengZichao/PyiTOL and archived with test data at Zenodo (https://doi.org/10.5281/zenodo.22106806).

## 1. Introduction

Phylogenetic trees organize hypotheses of evolutionary relationships and are routinely linked to taxonomic, functional and ecological metadata. Metagenomic and whole-genome studies can contain thousands of sampled taxa and multiple annotation layers [1–3]. At this scale, consistently generating, validating and versioning tree annotations becomes difficult, motivating programmatic workflows whose inputs, parameters and outputs can be reproduced.

The Interactive Tree of Life (iTOL) [4,5] is a widely used online platform for displaying and annotating phylogenetic trees with color strips, heatmaps, charts, symbols and tree-structure controls. Its annotation datasets are tab-delimited text files with type-specific identifiers, parameters and column layouts. When these files are assembled manually, errors in headers, delimiters, column order or required values can prevent import; repeated preparation across datasets also complicates batch processing and provenance tracking.

Existing software addresses separate parts of this workflow. At the versions evaluated in this study, itolapi [6] provides Python access to iTOL upload and export operations, whereas itol.toolkit [7] and table2itol [8] generate annotation files in R. Toytree [9] and ggtree [10] provide programmatic tree visualization rather than iTOL-template workflows. ETE3 [11] and ETE4 [12] classify annotated tip sets as monophyletic, paraphyletic or polyphyletic, while DendroPy [13] provides general tree operations, including MRCA queries, from which such tests can be constructed. However, no existing tool at the versions evaluated combines these capabilities into a single versioned Python command-line workflow.

To address these gaps, we developed PyiTOL (Python for iTOL), a command-line package that integrates validated annotation-file generation, rooted-tree taxonomic diagnostics, iTOL batch operations and provenance capture. Its methodological components are an explicitly defined, root-dependent mixed-child pattern classifier; separate reporting of sampling completeness, data adequacy and topology; and a deterministic post-order procedure that partitions a flagged member set into maximal member-only clades in O(n) time per group, where n is the number of tree nodes. We use the terms para- and polyphyly in the historical sense of Hennig [14] and Farris [15], but the present classification is operational and tree-dependent rather than a formal equivalence claim or a revised biological definition. These outputs therefore serve as auditable diagnostics for a specified tree and annotation mapping, not as new phylogenetic theory. The following sections state the assumptions, classification rule, pseudocode, complexity, software implementation and validation design.

## 2. Methods

### 2.1 Workflow and software architecture

PyiTOL uses a layered workflow that converts a phylogenetic tree and tabular annotations into validated iTOL datasets, optional taxonomic diagnostics, remote-service outputs and a provenance record (Figure 1). Newick and Nexus inputs are parsed before tip identifiers are reconciled with the annotation table. Ancestor-based analyses interpret each tree exactly as supplied: PyiTOL neither infers nor alters the root, so users should provide explicitly rooted trees; root-dependent results are reported relative to the supplied root (§2.3). Duplicate tip labels are rejected by the validator as hard errors, because set-based diagnostics cannot distinguish same-labelled leaves. Multi-tree files are handled by four named strategies (ask, first, last, split) plus explicit --tree-index selection, with automatic fallback to first in non-interactive environments. Configuration files are merged in their stated order, explicit command-line values take precedence, and the resolved configuration is written to the session record. Core routines perform parsing, validation and tree analysis; registered schemas generate annotation files; and a separate client handles authenticated iTOL operations. The command-line interface is implemented with Typer [16]; command-level details and the module map are provided in Supplementary Section S2.

**Figure 1.**
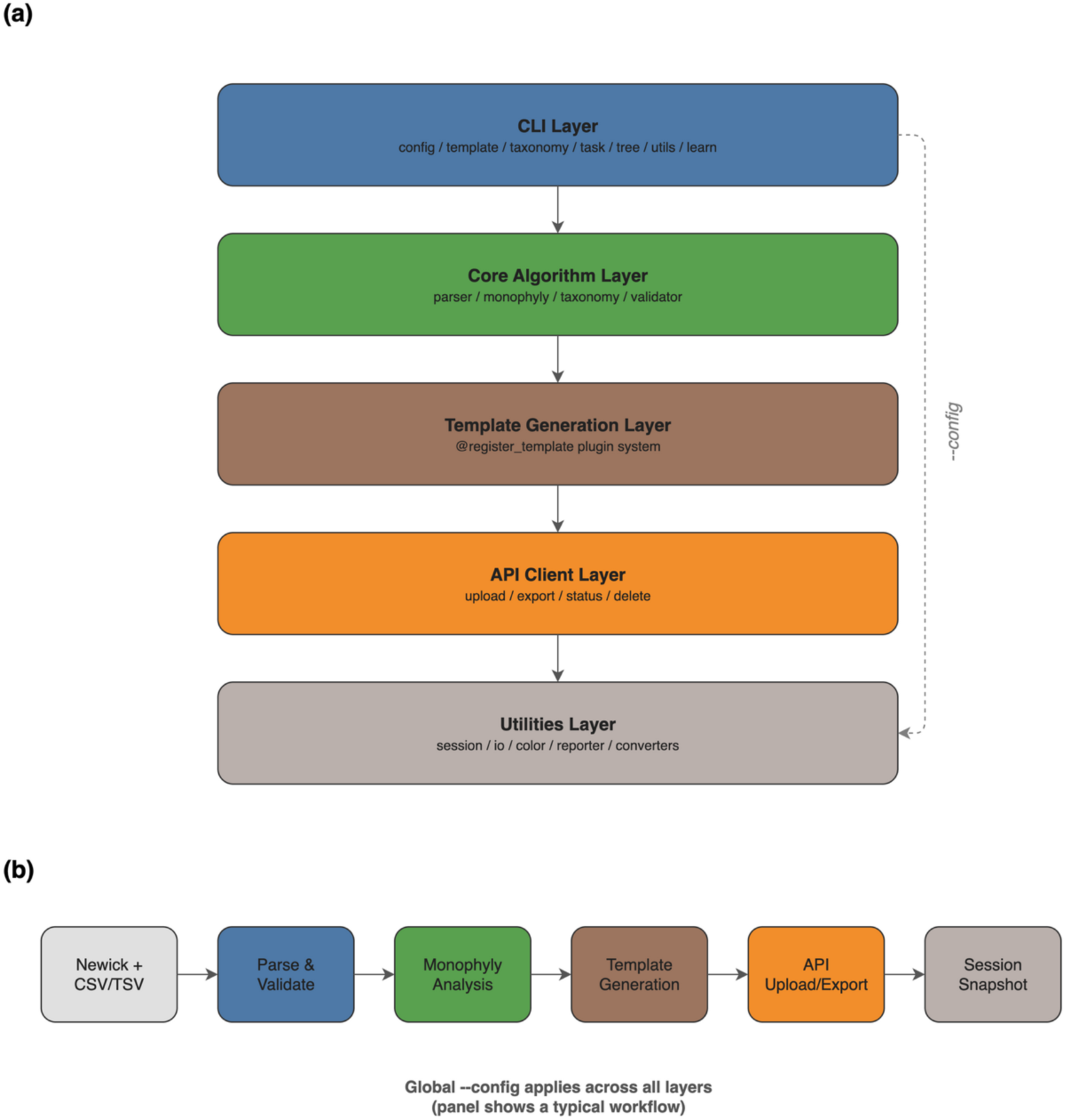
PyiTOL software architecture and workflow. Five-layer architecture (command-line interface, core algorithm, template generation, API client, utility) with left-to-right data flow from Newick/CSV input through parsing, validation, monophyly analysis, template generation and API upload to the session snapshot. Alt text: A five-layer software architecture diagram showing data flowing from Newick tree and CSV taxonomy inputs through command-line interface, core algorithm, template generation, and API client layers to iTOL upload, with a session snapshot recording the workflow.

### 2.2 Benchmark environment and reproducibility

All development and benchmark measurements were run on a single MacBook Pro with an Apple M5 processor and 32 GiB unified memory, using macOS 26.6.2 and Python 3.11.15 in a micromamba environment, and the automated test suite additionally runs on GitHub Actions CI (§4.1). The archived PyiTOL 1.0.3 release contains the dependency specifications (environment.yml and pyproject.toml), benchmark scripts and raw outputs (the reported benchmarks were run on v1.0.2; v1.0.3 differs only in input-validation strictness and retry logging). Supplementary Section S1 specifies the single-threaded execution settings, repetition counts, warm-up procedure, timed code boundaries and memory-measurement method; operating-system compatibility tests are reported separately from performance measurements.

### 2.3 Operational classification of taxonomic tip-set patterns

Let T be a tree interpreted as rooted, with a set U of unique tip identifiers, and let S be the expected identifiers assigned to taxonomic group G. Define the located members P = S ∩ U and the missing members M = S \ U; throughout, E and M are sets of tip identifiers. If P = ∅, no ancestor can be evaluated and the classifier reports unknown. Otherwise, let v = LCA(P), let L be the set of tip identifiers descended from v, and define the extra tips E = L \ P. Thus M records expected members absent from T (sampling absence, not topological scatter), whereas E records observed tips below v that are not located members; the topology call depends only on how E is distributed among the child clades of v. For a child c of v, let D(c) denote its descendant-tip set; c is mixed when D(c) contains both located members and extra tips. When tip labels are duplicated the validator reports a hard error before classification, because set operations would otherwise collapse distinct leaves. The classification is formalized as (Figure 2):

- **Monophyletic:** E = ∅ and M = ∅; or E ≠ ∅ but every extra tip maps back to G through the tip-to-group annotation (nested monophyly — e.g., extra tips belonging to genera contained in a checked family do not break its monophyly). This mapping rule is applied only at the queried rank and only when the annotation defines a group for every relevant tip; otherwise the classifier reports data_insufficient.
- **Paraphyletic:** M = ∅, E ≠ ∅, and the LCA has at most one mixed child — either S equals L minus whole sister clades (0 mixed children, the strict classical shape, reachable only at an unresolved LCA polytomy under relaxed handling, since strictly binary trees always yield at least one mixed child when E ≠ ∅) or all extra tips are nested inside the single subclade that also contains the remaining members (1 mixed child).
- **Polyphyletic:** M = ∅, E ≠ ∅, and the LCA has two or more mixed children — the located members are distributed across mutually disjoint subclades that each also contain extra tips. All labels in this section are operational and root-dependent: re-rooting the same unrooted topology can change the mixed-child count.

**Figure 2.**
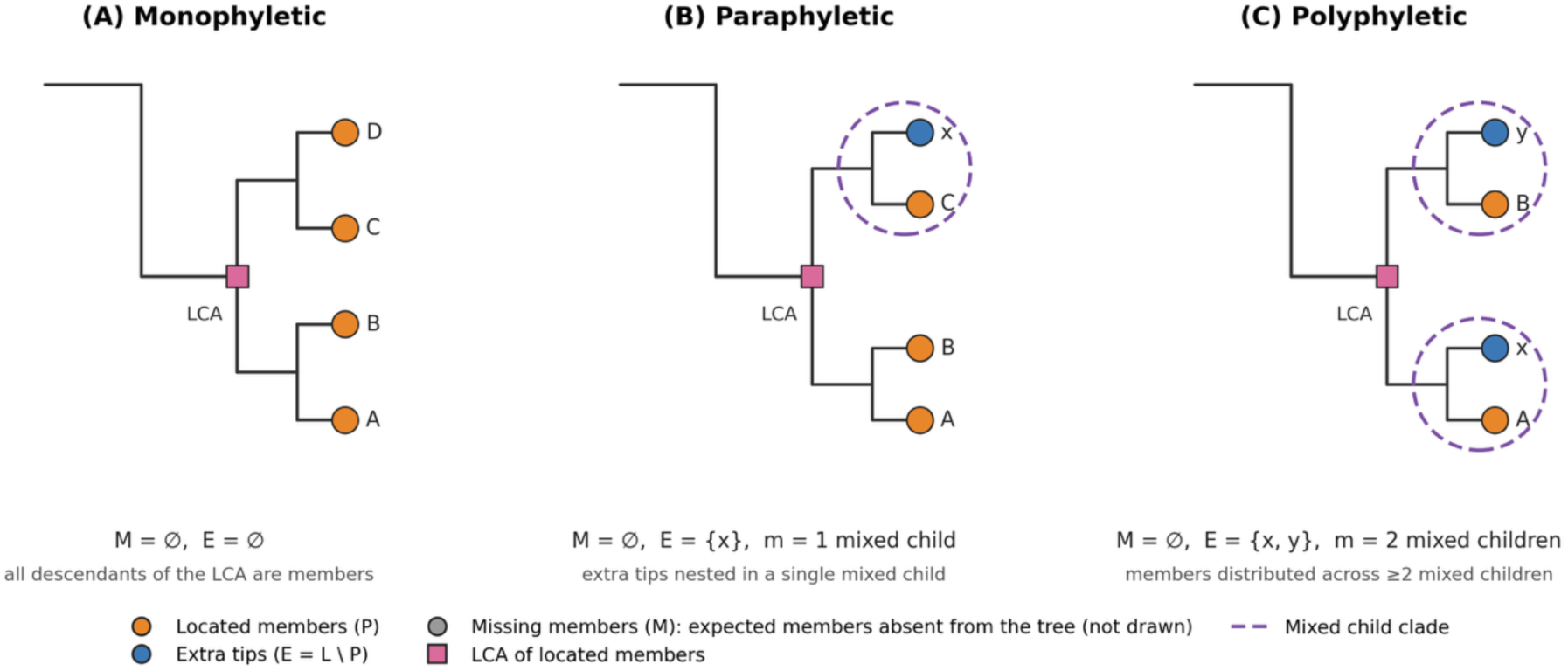
Schematic of the three-way classification of taxonomic tip-set patterns on a rooted tree. (A) Monophyletic: E = ∅ (and M = ∅); (B) paraphyletic: extra tips nested within at most one mixed child of the LCA; (C) polyphyletic: located members distributed across two or more mixed children. LCA positions and the E/M sets are annotated; missing members (M) are expected members absent from the tree and are therefore not drawn (M = ∅ in all three panels). Alt text: Three schematic rooted phylogenetic trees illustrating monophyletic, paraphyletic and polyphyletic distributions of located members (P), with the least common ancestor marked and the extra-tip set (E) annotated in each panel; the polyphyletic panel shows at least two mixed children; missing members (M) are absent from the tree and not drawn.

The mixed-children criterion implements an operational reading of Farris’s (1974) definitions [15]; it is a software decision rule, and no formal equivalence with the historical definitions is claimed:

**Proposition 1.** *Let M = ∅, E ≠ ∅, v = LCA(S) with descendant leaf set L, and let m(S) be the number of mixed children of v. Then: (a) m(S) = 0 if and only if E is a union of whole child clades of v — the strict classical shape (an ancestral descendant set minus entire sister clades); on strictly binary trees this shape is impossible when E ≠ ∅, so it is reachable only at multifurcating LCAs under relaxed polytomy handling. (b) If E is the complete descendant set of a single node — the paraphyly criterion implemented by ete3/ete4 — then m(S) ≤ 1; hence the ete3/ete4 paraphyletic class is contained in PyiTOL’s, and the inclusion is strict (witness: T = (A,((B,X),(C,Y))) with S = {A, B, C}: the single mixed child gives m = 1 and PyiTOL reports paraphyletic, whereas {X, Y} is the complete descendant set of no single node, so ete3/ete4 report polyphyletic). (c) m(S) ≥ 2 if and only if members of S occur in at least two child subclades of v that each also contain non-member tips; the classifier then reports polyphyletic under the operational definitions of §2.3.*

*Proof sketch. (a) If m = 0, every child of v is all-member or all-extra, so E is exactly the union of all-extra child clades; the converse is immediate; the binary-tree claim follows because v = LCA(S) forces members into at least two child clades or into one, and the latter contradicts the minimality of v. (b) Write E = Desc(z); since z ≠ v (as P ≠ ∅), all extra tips lie below a unique child of v, giving m ≤ 1; the displayed counterexample shows that the converse fails, hence the inclusion is strict. (c) Each mixed child places members and non-members in the same child clade, which is precisely m ≥ 2. ∎*

On strictly binary trees a paraphyletic group has exactly one mixed child (the m = 0 case requires a polytomy at the LCA), so Proposition 1 covers every paraphyly shape on resolved trees. Three supplementary states — incomplete_sampling when M ≠ ∅, data_insufficient when E contains unmapped tips or the LCA is an unresolved polytomy occupied by only a subset of member branches, and unknown when no member is located — are reported through a single precedence rule (unknown, then incomplete_sampling, then data_insufficient, then the topology label), so exactly one status is returned per group; the extra and missing tip sets remain available as additional output fields for inspection. Sampling completeness is evaluated before topology because absent members render any topology-based call unreliable. Unresolved LCA polytomies are handled by user-selectable modes: strict (default) returns data_insufficient, and data-insufficient is currently an identical alias retained for interface compatibility; relaxed is explicitly a heuristic that outputs such groups as paraphyletic for downstream review rather than as resolved topology (Supplementary Section S3). Formalization, pseudocode and complexity analysis are given in Supplementary Section S3.

## 3. Implementation

### Template generation engine

Each dataset type declares its metadata through a @register_template decorator executed at import time, populating SCHEMA_MAP, TEMPLATE_TYPE_HEADER and REQUIRED_COLUMNS; the factory create_schema(type_name) instantiates the corresponding schema, which TemplateGenerator serializes into iTOL v7-format text (Figure 3). Compatibility is empirically validated only for the 22 batch-validated types (§4.4); the remaining registered types are experimental pending the upgrades listed in §5.2. PyiTOL registers 31 template types — 27 dataset types plus 4 tree-structure types (COLLAPSE, PRUNE, SPACING, TREE_STYLE) — with the complete inventory in Supplementary Table S9. A _DEFAULT_LABELS mapping is verified by tests to give identical output from the unified entry pyitol template create <type> and the dedicated subcommands, and an LRU cache keyed by SHA-256 digests of the input paths, modification times and the taxonomy column parameter (invalidated by file modification time) avoids repeated parsing when generating multiple templates from the same input.

**Figure 3.**
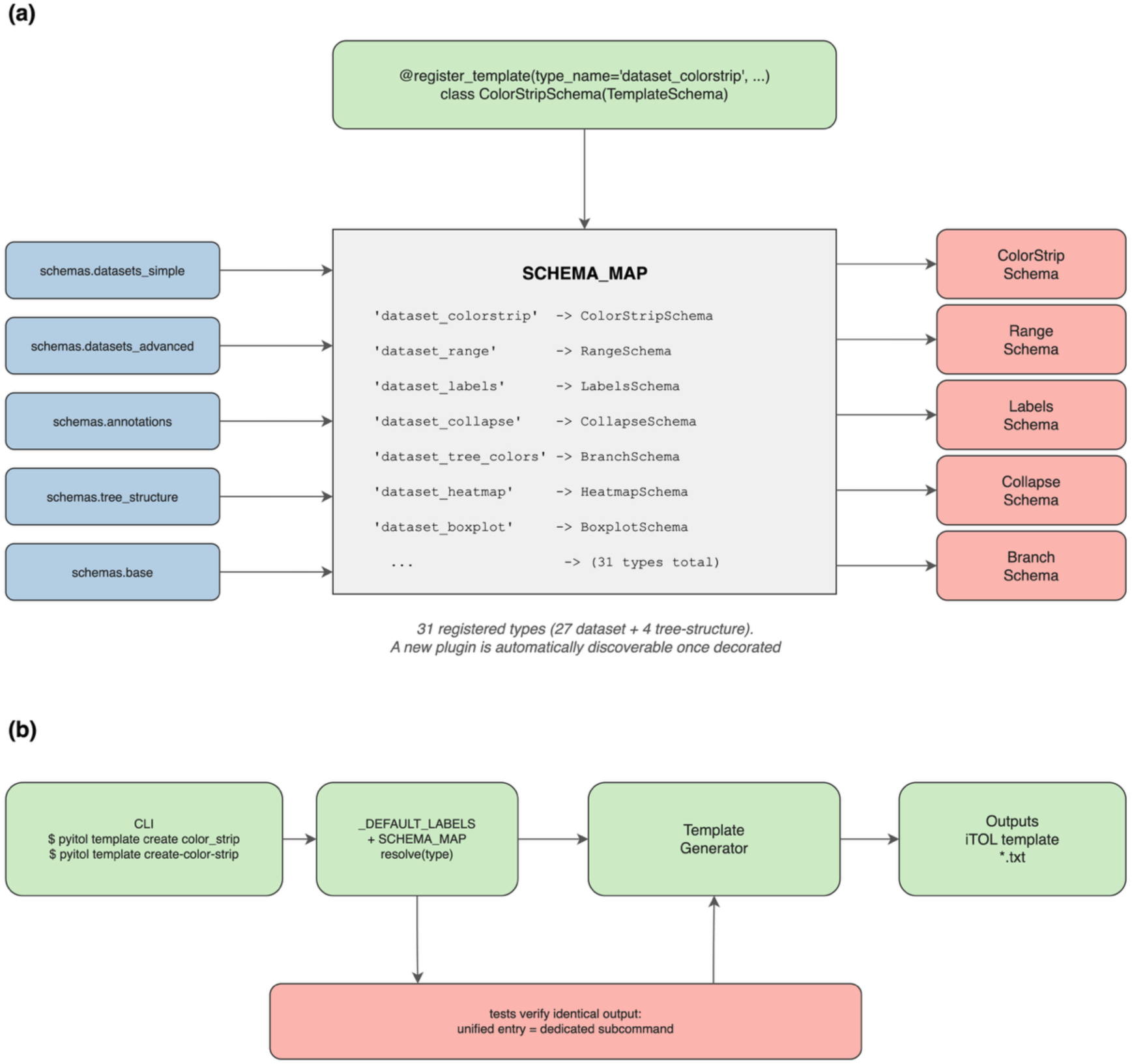
Template plugin registration system and unified-entry consistency. Left: the @register_template decorator populates `SCHEMA_MAP` at import time. Right: the unified entry uses `_DEFAULT_LABELS` to match dedicated subcommand defaults. Alt text: A diagram of the template plugin system: the register_template decorator populates a central schema map at import time, and a unified command entry selects type-specific default labels to match dedicated subcommands.

### API client

ITOLAPIClient packages the tree and templates into a single ZIP (tree files forced to the .tree extension), parses multiple iTOL response formats, retries with exponential backoff (timeout 120 s; up to 3 retries on HTTP 429/500/502/503/504 via urllib3 Retry), polls rendering status, and validates exports by checking HTTP status and content type first and then applying minimum-size thresholds and magic-byte prefixes as warnings rather than content parsing (Figure 4). Because the iTOL batch endpoints expose no idempotency keys, retried non-idempotent POST requests may create duplicate resources; the client therefore logs a warning on every retried POST, and users should verify the target project after any retried upload.

**Figure 4.**
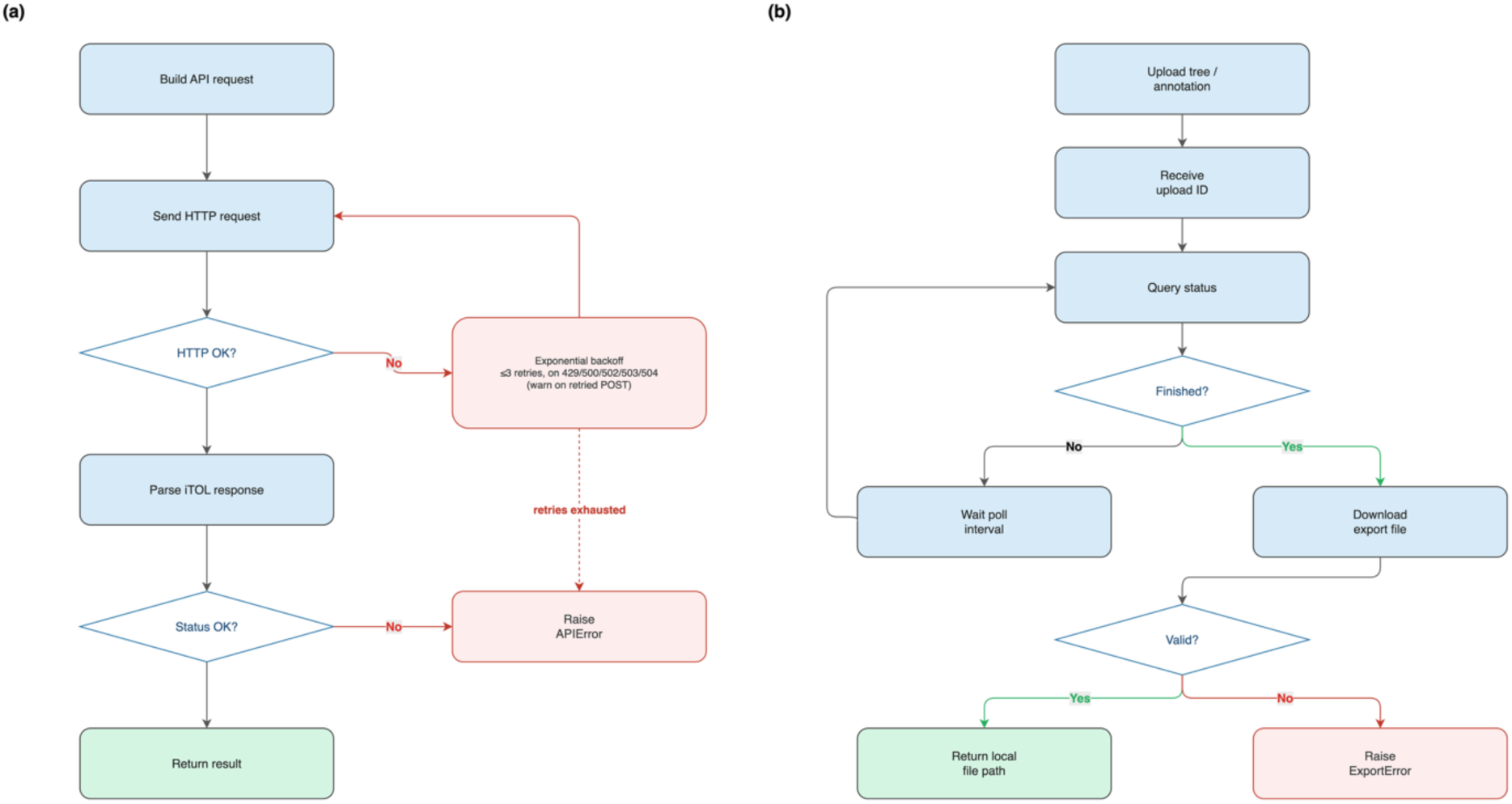
API-client retry, polling, and export-validation flow. (a) Generic request with exponential-backoff retry; (b) export flow with upload, status polling, download and file-validity checks. Alt text: Two flowcharts: (a) an API request path with exponential-backoff retry on server errors, and (b) an export path with upload, status polling, download and file-validity verification steps.

### Validation, robustness and reproducibility

safe_read_text() strips UTF-8 BOM prefixes (U+FEFF) prepended by some editors and falls back to utf-8-sig decoding; other encodings (e.g., UTF-16) are out of scope. load_tree() supports four named multi-tree strategies (ask, first, last, split) plus explicit --tree-index selection, with automatic fallback to first in non-interactive environments; the validator checks color codes, delimiter conflicts, numeric ranges and duplicate tip labels (hard errors); signal-safe interruption supports clean Ctrl+C exit (implementation details and robustness test results in Supplementary Section S8). SessionContext serializes session snapshots (session ID, software versions, command line, input SHA-256 checksums, API parameters with keys automatically redacted) validated by a Pydantic model [17]; pyitol replay re-executes the recorded local steps from a snapshot; bit-identical remote results are not guaranteed, as external services are not snapshotted. Errors follow a structured exception hierarchy with an internationalization-ready message catalog currently shipping English and Chinese templates (PYITOL_LANG).

## 4. Results

### 4.1 Software quality and comparison with existing tools

The automated test suite comprises 72 modules with 1,704 test cases covering parsing, validation, monophyly analysis, template generation, the API client, configuration and session management; CI runs on GitHub Actions at 86.5% statement coverage with a --cov-fail-under=80 floor (Supplementary Sections S4 and S6, Supplementary Figure S1). Table 1 compares PyiTOL functionally with existing tools; all capability judgments were verified against each tool’s source code and documentation at the tested versions (itolapi 4.1.6, itol.toolkit 1.2.2, table2itol commit of July 2026, ete3 3.1.3, ete4 4.4.0, DendroPy 4.6.4). The visual feature matrix is given in Supplementary Figure S2.

**Table 1.**
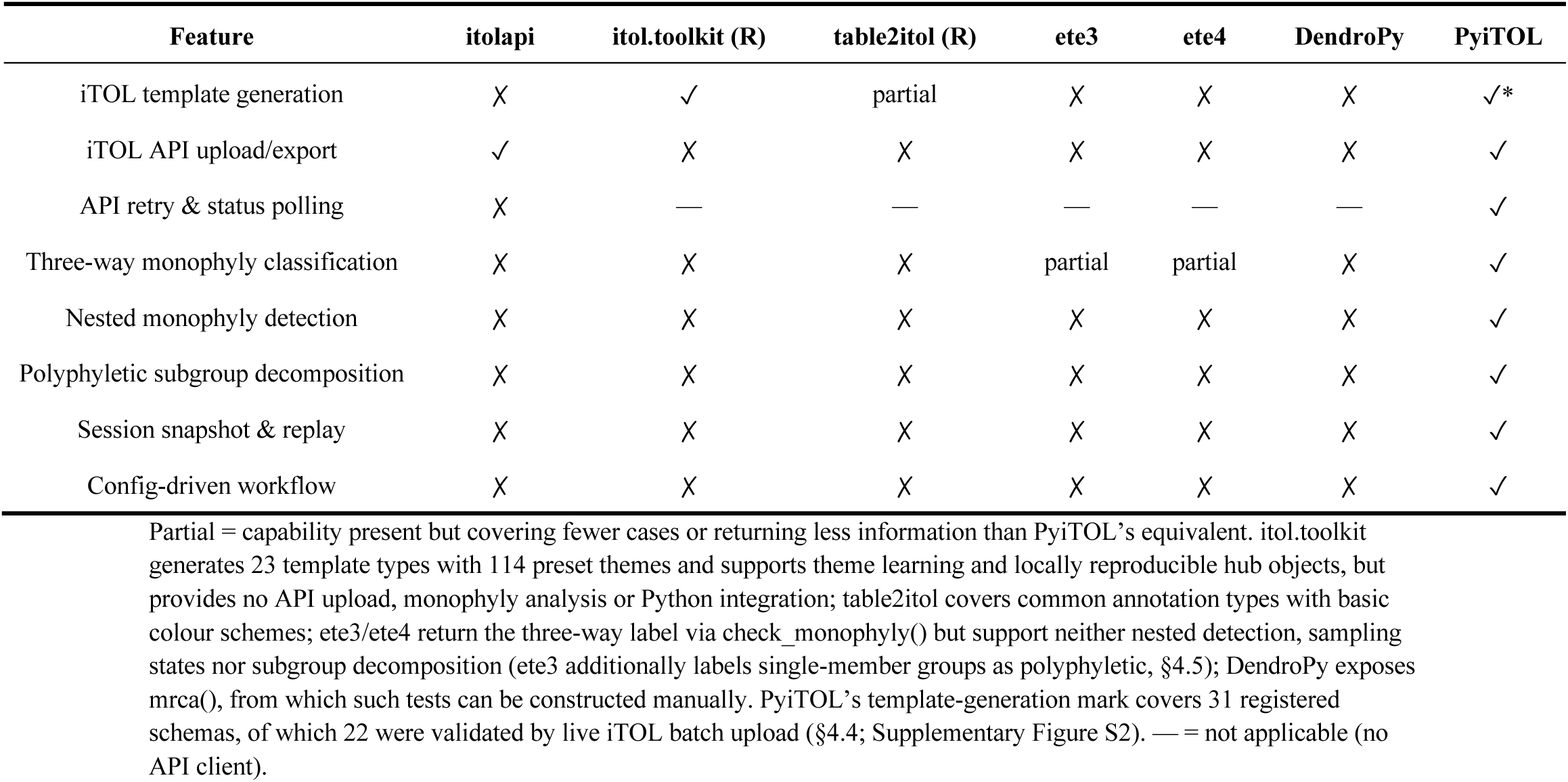
Functional comparison of PyiTOL with existing tools.

### 4.2 Rule conformance on a constructed benchmark

To quantify rule conformance at scale we built a simulation benchmark (an implementation-conformance suite rather than a test of biological accuracy) in which ground truth is defined by topology construction (hence non-circular), covering two regimes: deterministic fully balanced binary trees (2,048 and 8,192 leaves; 5 seeds each) and seeded random-bifurcating (unbalanced) trees (2,048 and 4,096 leaves; 3 seeds each). Six categories were engineered per replicate — monophyletic, paraphyletic (including a deeply nested variant), polyphyletic, incomplete-sampling, data-insufficient and whole-clade filler groups — and every replicate was scored end-to-end through the pyitol taxonomy monophyly CLI in strict mode. Eleven additional boundary scenarios (polytomies under all three modes, single-member and whole-tree groups, nested monophyly, unmapped extras, empty and missing members) passed as expected (Supplementary Table S2).

Across 4,389 scored groups, all calls matched the prespecified constructed labels in both regimes (4,389/4,389; all off-diagonal entries of the merged confusion matrix are zero; Supplementary Table S4), and for all 397 engineered polyphyletic groups the subgroup decomposition recovered exactly the two expected maximal monophyletic subgroups. Because the engineered ground truth adopts PyiTOL’s own classification criterion, these scores quantify implementation fidelity to that criterion — they are not estimates of biological accuracy or of robustness to annotation error, and the benchmark additionally served as a regression suite during development, so conformance may be optimistic for unseen scenarios; subgroup diversity is also limited, as every engineered polyphyletic case expects exactly two subgroups. Behaviour on noisy empirical trees additionally depends on the polytomy-mode choice (§2.3) and upstream collapsing of low-support branches (§5.2); the external behaviour of the criterion on real data is examined in §4.5. (Table 2)

**Table 2.**
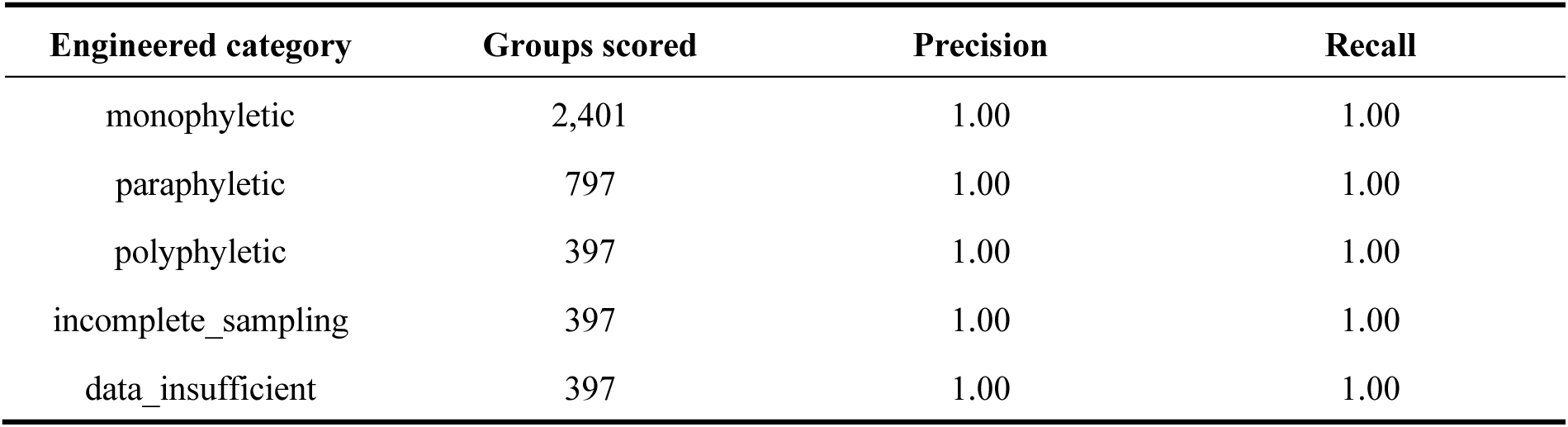
Simulation-based rule conformance (topology-constructed labels)

### 4.3 Computational efficiency

Monophyly determination was benchmarked against ete3, ete4 and DendroPy [13] on deterministic balanced binary trees (3 replicates per size; medians; identical input trees and taxonomies; all competitor versions pinned as listed in Supplementary Section S1). PyiTOL’s timed path is the conservative end-to-end scope (tree and taxonomy re-parsed per repeat); the matched-scope variant without parsing is 19.0 s at 50,000 leaves (Supplementary Table S6), within the same order of magnitude as ete3. (Tables 3 and 4)

**Table 3.** Performance comparison of monophyly determination: median wall-clock time across configurations (PyiTOL reports the full 10-field output; control tools return a single Boolean/trichotomy label; PyiTOL’s timed path re-parses inputs per repeat while the DendroPy control reuses a pre-parsed tree — matched-scope variants in Supplementary Table S6)

| Leaves | PyiTOL | ete3 | ete4 | DendroPy |
| --- | --- | --- | --- | --- |
| 1,000 | 0.41 s | 0.26 s | 0.09 s | 0.06 s |
| 10,000 | 4.74 s | 3.44 s | 0.96 s | 2.33 s |
| 50,000 | 23.47 s | 18.60 s | 4.84 s | 63.99 s |

**Table 4.** Performance comparison of monophyly determination: peak memory across configurations (tracemalloc, Python-level allocation; C/Cython allocations may be underestimated, so cross-language rankings are indicative only; PyiTOL reports the full 10-field output while control tools return a single Boolean/trichotomy label)

| Leaves | PyiTOL (mem) | ete3 (mem) | ete4 (mem) | DendroPy (mem) |
| --- | --- | --- | --- | --- |
| 1,000 | 2.5 MiB | 2.4 MiB | 738 KiB | 76 KiB |
| 10,000 | 35.8 MiB | 27.8 MiB | 7.9 MiB | 1.1 MiB |
| 50,000 | 439.0 MiB | 139.1 MiB | 37.5 MiB | 4.5 MiB |

For the k-group batch, the timed competitor runs perform a per-query label scan for each of the k group comparisons, precisely the cost eliminated by PyiTOL’s _tip_index_cache, which reduces repeated member localization from O(n·k) to O(n + k). Peak memory was measured with tracemalloc (Python-level allocation; C-extension allocations may be underestimated, so cross-language memory rankings are indicative only). On stress workloads in which nearly all taxa are polyphyletic, 100,000- and 200,000-leaf synthetic balanced trees required 212.0 s (206.6–212.3 s across three replicates; 1.6 GiB) and 424.2 s (422.6–424.7 s; 5.7 GiB), respectively (Table 5; reproduction script in Supplementary Section S5).

**Table 5.** Extreme-scale stress benchmark on synthetic balanced binary trees.

| Dataset | Leaves | Groups checked | Wall time (median, min–max) | Peak memory |
| --- | --- | --- | --- | --- |
| Synthetic balanced binary tree | 100,000 | 100 | 212.04 s (206.61–212.28 s) | 1.6 GiB |
| Synthetic balanced binary tree | 200,000 | 100 | 424.22 s (422.63–424.74 s) | 5.7 GiB |

Template-generation time was approximately proportional to tree size over the measured range: color-strip, heatmap and simple-bar templates for a 50,000-leaf tree complete in about 2.0 s with peak allocations of 92.0/14.7/13.6 MiB (runtime and memory also depend on the number of data columns, categories and output size; Supplementary Table S7).

### 4.4 Live-server acceptance of generated templates

For each of the 31 registered types, a minimal valid template was generated against a 32-tip example tree and uploaded to the live iTOL server (v7.6; tested 2026-08-04) through PyiTOL’s API client; acceptance required SUCCESS with a tree ID, which confirms tree upload but not dataset attachment or rendering — accepted types were additionally inspected visually in the iTOL web interface. 22 of 31 types (71%) were accepted on first submission (Table 6), covering all high-frequency annotation workflows. The nine non-accepted types fall into distinct cause categories rather than a single server-side limit: (i) DATASET_IMAGE is documented by iTOL as unsupported by the batch uploader; (ii) DATASET_MANUAL and DATASET_PLACEMENT have no tabular-template equivalent in iTOL’s own workflow (manual is an interactive drawing layer; placement data are derived from .jplace files); (iii) DATASET_TANGLEGRAM is currently generated as the legacy tabular id1/id2 mapping, which does not cover the iTOL v7 structure requiring a second tree embedded between TANGLEGRAM_TREE/END_TANGLEGRAM_TREE — a generator upgrade listed as future work (§5.2); and (iv) PRUNE, DATASET_TIMESCALE, DATASET_TREESTYLE, DATASET_ARROWS and DATASET_MEME are applied through iTOL’s web interface or control panel and are not identified by the batch uploader. All nine types remain generable for manual upload through the web interface; per-type server responses and interpretation are given in Supplementary Table S3, and raw responses are archived with the repository (benchmarks/itol_acceptance/acceptance_results.json), allowing readers without an iTOL subscription to inspect the evidence behind each call.

**Table 6.** Live iTOL server acceptance of PyiTOL-generated templates.

| Outcome | Count | Template types |
| --- | --- | --- |
| Accepted (SUCCESS with tree ID) | 22 | alignment, binary, boxplot, branch/tree_colors, collapse, color_strip, connection, domains, externalshape, gradient, heatmap, labels, linechart, multi_bar, pie, popup_info, range, simple_bar, spacing, style, symbol, text |
| Rejected: explicit batch-mode limitation | 1 | image |
| Rejected: type not identified by the batch uploader (failure causes (i)–(iv) in §4.4) | 8 | arrows, manual, meme, placement, prune, tanglegram, timescale, treestyle |

### 4.5 Real biological case studies

#### GTDB R232 reference trees

We used the two reference trees released with GTDB R232 [18,36]: ar53 (53 archaeal marker genes, 10,122 genomes) and bac120 (120 bacterial marker genes, 189,801 genomes), redistributed with the archived repository under data/gtdb_r232/ (original data at https://gtdb.ecogenomic.org/downloads). Terminal taxonomy was derived from internal-node labels by scripts/extract_gtdb_taxonomy.py (reproduction commands in Supplementary Section S7); suffixed taxa (e.g., Bacillota_I) were treated as distinct taxa throughout, following GTDB’s rank-tree convention. Phylum-level color-strip templates were generated in about 0.7 s (ar53) and 8 s (bac120); genus-level monophyly validation completed in 0.95 s for ar53 (1,322 genera) and about 17 s for bac120 (17,294 genera; peak memory about 4.9 GiB, Supplementary Section S7.2). The genus-level evaluation covered genera with at least two member genomes, running with --polytomy-mode relaxed; all evaluated genera were classified as monophyletic. Because GTDB genera are curated to be monophyletic and the terminal taxonomy used here is extracted from the internal-node labels of the same trees under test, this agreement demonstrates pipeline consistency and scaling behaviour; it cannot establish classifier accuracy or the absence of false positives. True-positive assessment is therefore carried out separately below. Collapsed reference trees with genome-quality indicators are shown in Figure 5.

**Figure 5.**
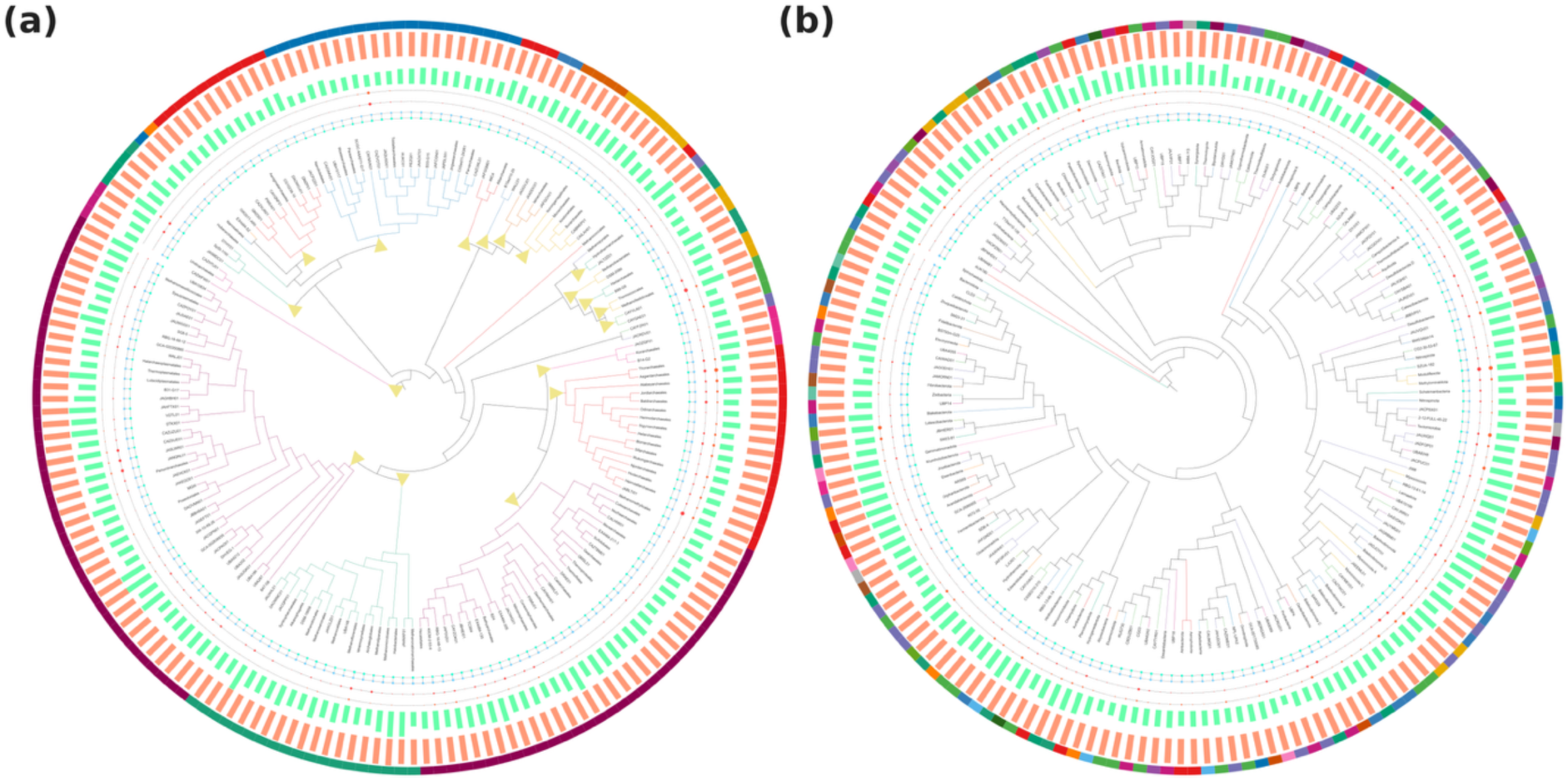
GTDB R232 reference trees collapsed and annotated with PyiTOL. (A) Archaeal ar53 tree collapsed at order level (179 terminals); (B) bacterial bac120 tree collapsed at phylum level (172 terminals). Annotation layers, from the tree outward: phylum-coloured branches (fixed palette keyed by phylum initial, with identical mapping in both panels), MRCA markers for collapsed clades (panel A only), genome completeness and contamination bubble tracks, GC-content and genome-size bar tracks, and an outer phylum colour strip. Suffixed GTDB taxa (e.g., Bacillota_I) are treated as distinct taxa throughout. Numeric legends are embedded in the corresponding iTOL template files; full colour mappings and rendering parameters are given in Supplementary Section S7. Genome-quality indicators (completeness and contamination per CheckM and CheckM2, GC content, genome size) originate from the GTDB R232 metadata table redistributed with the repository; values shown for collapsed clades are arithmetic means over member genomes, with genomes missing a given metric excluded from that mean; per-layer inputs and provenance are listed in Supplementary Section S7. Alt text: Two circular phylogenetic trees of GTDB R232 archaeal and bacterial reference trees collapsed at order and phylum level, surrounded by annotation layers for phylum colours, collapsed-clade markers, genome completeness and contamination indicators, GC content and genome size.

#### Detection of non-monophyletic taxa on a published phylogenomic tree

Because the GTDB case cannot demonstrate true positives, we additionally evaluated PyiTOL on a published relaxed molecular-clock phylogenomic tree of 700 archaeal and bacterial genomes [19], obtained from the publicly archived figshare dataset [20] (CC BY 4.0). The file was renamed and its tip labels re-annotated by the present authors to embed six-rank taxonomy; the topology is unmodified (provenance in Supplementary Section S7). Batch checks at every rank in strict mode flagged candidate non-monophyletic taxa relative to the supplied rooted topology and the re-annotated taxonomy (Table 7).

**Table 7.** Candidate non-monophyletic taxa on the 700-genome phylogenomic tree (tree-relative calls under the supplied rooted topology and re-annotated taxonomy; at genus level, 386 of 409 taxa are single-member groups, so the informative comparisons are the 23 multi-member genera)

| Rank | Taxa checked | Monophyletic | Paraphyletic | Polyphyletic | Data insufficient |
| --- | --- | --- | --- | --- | --- |
| genus | 409 | 406 | 2 | 1 | 0 |
| family | 231 | 209 | 14 | 6 | 2 |
| order | 155 | 133 | 10 | 7 | 5 |
| class | 88 | 75 | 8 | 4 | 1 |
| phylum | 125 | 119 | 4 | 2 | 0 |
| domain | 2 | 2 | 0 | 0 | 0 |

At genus level, exactly three non-monophyletic genera were flagged, all with clear topological explanations: Sulfolobus was polyphyletic — its two genomes fall into two distant Sulfolobaceae lineages and were decomposed into exactly two maximal monophyletic subgroups, each containing one of the two genomes (a weak-evidence candidate consistent with the ongoing phylogenomic splitting of this genus); Archaeoglobus was paraphyletic with Ferroglobus and Geoglobus nested inside its LCA clade (2 extra tips, single mixed child); and Thermococcus was paraphyletic with Palaeococcus nested between two Thermococcus genomes (1 extra tip). Because 386 of the 409 genus-level taxa are single-member groups whose monophyly call is trivially true, the informative comparisons are concentrated in the 23 multi-member genera. All 409 genus-level calls were cross-validated against ete3 (3.1.3) and ete4 (4.4.0) [11,12] on the same tree (Supplementary Table S5): PyiTOL and ete4 agreed on 409/409 binary (monophyletic/non-monophyletic) calls. ete3 agreed on every multi-member group but labels all 386 single-member genera polyphyletic and classified Thermococcus as polyphyletic; both quirks trace to ete3’s get_common_ancestor() falling back to the tree root when queried with a single node. These results are tree-relative candidates that require external taxonomic evidence, branch support and topology sensitivity analyses before biological confirmation; they nonetheless illustrate the practical value of the explicit three-way-plus-states semantics.

## 5. Discussion

### 5.1 Contributions

PyiTOL’s primary contribution is integrating the end-to-end iTOL annotation workflow — data validation, taxonomy analysis, plugin-based generation of 31 template types (22 batch-validated; local generation plus remote upload, with acceptance boundaries quantified in §4.4), API upload and session replay — into a single Python CLI, filling a gap in the Python ecosystem where no equivalent of the R-based itol.toolkit existed. Second, it turns an operational reading of the classical Hennig–Farris three-way monophyly framework (§2.3) into a batch-executable validation tool, extended with nested-monophyly detection, sampling-completeness states and exact maximal-clade subgroup decomposition; the mixed-children criterion is formally characterized (Proposition 1), and the simulation evaluation verified implementation conformance of the classifier and exposed — and led us to replace — a defective greedy subgroup implementation with an exact O(n)-per-group algorithm. Third, live-server acceptance testing quantified what the iTOL batch uploader accepts from generated templates (22/31) and classified the nine non-accepted types by failure cause (§4.4). Fourth, session snapshots with input checksums and redacted API parameters provide auditable reproducibility for local processing, while iTOL’s server-side rendering remains outside snapshot control by design. The internationalization-ready message catalog (English/Chinese) lowers the barrier for non-English-speaking users.

### 5.2 Limitations and future work

The polytomy-handling boundary and its operating modes are formalized in Supplementary Section S3. The strict default returns data_insufficient for unresolved LCA polytomies (data-insufficient is currently an identical alias retained for interface compatibility), while relaxed is explicitly a heuristic: it outputs such groups as paraphyletic for downstream review in taxonomic-revision workflows [21], not as a determination that the topology is resolved. Paraphyly/polyphyly discrimination additionally assumes that child-clade structure is resolved; for noisy trees we recommend first inferring trees with bootstrap support using RAxML [22], IQ-TREE [23] or FastTree [24] (or phylogenomic placement pipelines such as PhyloPhlAn [25]), collapsing low-support branches, and then running relaxed mode, unifying terminal labels with pyitol validate before checking, combining MAG-tree results with CheckM [26] quality scores, with taxonomic inputs commonly derived from GTDB-Tk [27,28] or 16S rRNA databases such as SILVA [29]. The deployment thresholds used in Supplementary Section S7 (e.g., bootstrap < 50 for collapsing, ∼100,000 leaves for partitioning) are empirical heuristics, not universal rules: appropriate cut-offs vary with data, model and hardware, and partitioning trees by phylum/class can hide non-monophyly that spans partition boundaries; sensitivity checks are advised. Full deployment guidance is given in Supplementary Section S7.

Current limitations are as follows. iTOL batch upload requires an active subscription and a pre-created project; offline mock tests keep CI green, but real-server tests depend on user credentials, and iTOL’s lack of a formal API specification means response-format changes could require parsing updates. Because the batch endpoints expose no idempotency keys, retried POST uploads may create duplicate trees on the server; PyiTOL logs a warning on every retried POST and users should check the target project afterwards. PyiTOL supports Newick and Nexus but not PhyloXML [30] or NeXML [31], and complex Nexus annotations (e.g., BEAST [32] output) may lose metadata. All template generators follow a data-driven “taxonomy column → formatting” paradigm without topology-aware automation (e.g., clade colouring requires user-supplied MRCA ids), a boundary shared with itol.toolkit [7]. Template generation and monophyly analysis are single-threaded; the Tables 3–4 memory figures reflect the information-for-memory trade-off of the full 10-field output, and workstations with ≤16 GiB RAM should partition trees above ∼100,000 leaves. Types not yet batch-verifiable — tanglegram (embedded second tree) and placement (.jplace) — are currently unsupported in batch mode and are listed below as future work rather than verified capabilities. Future work includes --batch-size multithreading, polytomy-resolution metrics, a NeXML reader via DendroPy [33] and a PhyloXML reader via a dedicated parser, a topology-aware - -mode clade for TREE_COLORS, real-API CI via GitHub Actions secrets, batch-mode upgrades for tanglegram and placement types, and streaming template generation building on the existing --low-memory mode.

## Availability and Data Access

### Software availability

PyiTOL is open-sourced under the MIT license, which applies to the PyiTOL source code; redistributed GTDB and Moody et al. data remain under their respective licences (Data availability). The submission reviews version 1.0.3 (tag v1.0.3). Source code, test data, example datasets and benchmark artifacts are freely available at https://github.com/ZengZichao/PyiTOL and archived at Zenodo (https://doi.org/10.5281/zenodo.22106806) [34]; the package installs via pip install pyitol (Python ≥ 3.10; Linux, macOS, Windows). Remote iTOL operations additionally require an active iTOL subscription and API key. Documentation, tutorials and Jupyter notebooks [35] are hosted at https://zengzichao.github.io/PyiTOL. System requirements, Docker deployment and maintenance plans are given in Supplementary Section S2; the authors commit to maintaining availability for at least two years following publication.

### Data availability

The GTDB R232 reference trees are publicly distributed by GTDB (https://gtdb.ecogenomic.org/downloads; [18,36]) under the Creative Commons Attribution-ShareAlike 4.0 International (CC BY-SA 4.0) license, and are redistributed with the archived repository under data/gtdb_r232/ for convenience. All derived files (*_taxonomy.csv, *_phylum_strip.txt, *_monophyly_genus.csv) are regenerable with scripts/extract_gtdb_taxonomy.py and the commands in Supplementary Section S7; the redistributed copies and these derived files are therefore licensed under the same CC BY-SA 4.0 terms, with attribution to GTDB. The 700-genome phylogenomic tree originates from Moody et al. [19,20] (CC BY 4.0; redistributed with attribution, modifications documented in Supplementary Section S7).

## Author contributions (CRediT)

Z.Z.: Conceptualization, Software, Validation, Formal analysis, Investigation, Writing – original draft. Y.W.: Supervision, Project administration, Funding acquisition, Writing – review & editing.

## Use of AI assistance

During the preparation of this work the authors used an LLM-based assistant for (i) language editing and polishing of the manuscript text and (ii) assistance in writing source code, including algorithm implementations and helper functions. All scientific content was conceived, executed and verified by the authors, who reviewed and revised all AI-generated content and take full responsibility for the correctness and integrity of the code and the final manuscript. Detailed disclosure is given in Supplementary Section S10.

