## Supporting Information for "PyiTOL: reproducible Python workflows for iTOL annotation and taxonomic monophyly assessment"

### S1 | Benchmark Environment

All benchmarks in the main text were run on a single MacBook Pro (Apple M5 chip, 32 GiB unified memory) running macOS 26.6.2, with Python 3.11.15 managed by micromamba. Library versions: PyiTOL 1.0.2, DendroPy 4.6.4 (benchmarked version; the current v5.0 release line is cited as [33], as of August 2026), ete3 3.1.3, ete4 4.4.0, pandas 2.3.3, typer 0.26.8, numpy 2.4.6, scipy 1.17.1 (environment.yml and pyproject.toml declare the corresponding dependency ranges; the archived PyiTOL 1.0.2 release — tag v1.0.2, commit 8daec4c726d22fe194f515771436da560623599b — bundles the dependency specifications (environment.yml and pyproject.toml) used for these benchmarks; the reviewed submission version is PyiTOL 1.0.3 (tag v1.0.3, commit b9a120706e998cb9bf4f12c301c040ddcecd0222), which differs from v1.0.2 only in input-validation strictness and retry logging). No HPC cluster or cloud compute resources were used.

Template-generation benchmarks used benchmarks/benchmark_scalability.py. Synthetic trees were generated by DendroPy's Yule pure-birth process (birth_rate=1.0, death_rate=0.0) with tree_length=1.0 and ntax variable per benchmark size; trees exceeding numerical stability thresholds were generated via the build_balanced_newick() balanced binary tree fallback. A warm-up pass (all three template types on a 50-leaf tree) precedes all measurements so that lazy imports and first-call caches do not bias any template type; every reported memory value is therefore a tracemalloc incremental allocation measured under an identical methodology across template types. Each size was repeated 10 times, and median wall-clock time and peak memory are reported (per-replicate min/max ranges in benchmarks/template_scalability_results.json). Synthetic taxonomy columns are generated with a fixed seed (42); the Yule topology generator is not re-seeded across sizes, and trees exceeding the stability threshold switch to the balanced fallback — readers should therefore note that topology differences may contribute to size effects alongside leaf count (per-size inputs and all replicate values are recorded in the results JSON).

Monophyly horizontal-comparison benchmarks used deterministic fully balanced binary trees (recursive halving; `generate_exact_newick()` in `benchmarks/benchmark_monophyly_comparison.py`) at 1,000/10,000/50,000 leaves with 3 replicates per size, reporting medians. In addition to the end-to-end PyiTOL scope (which re-parses the tree and taxonomy from disk on every repeat) and the DendroPy scope (pre-parsed in-memory tree), a matched-scope PyiTOL variant (`pyitol_inmem`, full 10-field output on a pre-parsed tree, tip-index cache rebuilt inside each timed run) and a Boolean-only lightweight variant (`pyitol_light`, same MRCA + descendant-set logic as the DendroPy control) are reported in Table S6. In the DendroPy benchmark, `tree.is_rooted = True` was set before each `tree.mrca()` call to match PyiTOL's `_safe_mrca()` behaviour and eliminate rooted state as a confounding variable.

The cProfile hotspot analysis quoted in main-text §4.3 was obtained with Python's built-in `cProfile` module wrapping one full batch of the DendroPy control at 50,000 leaves (20 groups × 2,500 members, identical inputs to main-text Table 3); cumulative per-function inclusive times were read from the standard `pstats` output. The comparison taxonomy was generated by round-robin assignment (i % num_groups, num_groups = 20): each group contains n/20 members (2,500 at 50,000 leaves), and nearly all groups are polyphyletic, representing a stress scenario in which every call triggers the subgroup-decomposition path and the full 10-field output, whereas ete3, ete4 and DendroPy can early-exit after a single Boolean mismatch.

Runtime attribution note: parsing therefore accounts for only about 14–21% of PyiTOL's end-to-end runtime; the algorithmic determination itself (matched scope) remains within the same order of magnitude as ete3 (18.60 s at 50,000 leaves) while returning the full 10-field structured output. Under this stress workload, the full-output variant runs about 2.7× faster than the DendroPy control (63.99 s); because the two return different output widths (10 structured fields versus a single Boolean), this number characterises PyiTOL's full-mode throughput rather than an equal-task algorithmic advantage. The Boolean-equivalent variant (pyitol_light) runs at about 1.17× the DendroPy control at 50,000 leaves; the difference reflects library-internal overhead rather than algorithmic divergence (Table S6).

### S2 | Software Architecture and Deployment

#### S2.1 Module map

**Command-line interface layer** (built on Typer): seven command groups (`config`, `template`, `taxonomy`, `task`, `tree`, `utils`, `learn`) with subcommands, plus three top-level commands (`validate`, `replay`, `self-test`). The `learn` group learns reusable styling knowledge from previously published iTOL template files (extracting palette and parameter patterns into configuration files that can be re-applied during template generation). Command groups are registered via `app.add_typer()`; top-level commands are mounted via `@app.command()`. Typer's type-hint mechanism automates argument parsing, help generation and type validation. The global `--config` option can be specified multiple times; multiple YAML/JSON configuration files are merged in the order provided; explicit non-default CLI values take precedence over configuration files, which in turn take precedence over Typer defaults. Parameter aliases (e.g., `tree_path ↔ tree`, `taxonomy_path ↔ taxonomy`, `columns ↔ column`) lower migration cost.

**Core algorithm layer**: `core/parser.py` (Newick/Nexus auto-detection, BOM-safe reading, multi-tree processing), `core/monophyly.py` (three-way classification), `core/taxonomy.py` (embedded-label and GTDB-format extraction, rank member lists), `core/validator.py` (color codes, delimiter conflicts, numeric ranges, node-ID consistency), and data-conversion modules (`core/binary_convert.py`, `core/connect_convert.py`, `core/column_group.py`, `core/count_tree.py`).

**Template generation layer:** TemplateRegistry in templates/schemas/base.py provides centralized registration; the @register_template decorator writes type names, header identifiers and required columns into the registry at import time and exposes SCHEMA_MAP, TEMPLATE_TYPE_HEADER, REQUIRED_COLUMNS and NO_LABEL_COLOR_TYPES; create_schema(type_name) instantiates schemas; TemplateGenerator serializes them into iTOL v7-format text (batch acceptance verified for 22 of the 31 registered types; §S11 Table S3, main text §4.4).

**API client layer**: wraps iTOL batch upload, export, status query and delete endpoints with ZIP packaging, exponential-backoff retry, HTML error-page detection and polling-based rendering wait.

**Utility layer**: session context (`utils/session.py`), safe file reading and BOM handling (`utils/io.py`), color assignment (`utils/color_tools.py`), data conversion (`utils/data_converters.py`), FASTA handling (`utils/fasta.py`), tree information extraction (`utils/tree_info.py`), real-time logging (`utils/logger.py`) and bilingual error reporting (`utils/reporter.py`).

#### S2.2 System requirements

- Python ≥ 3.10
- Operating systems: Linux, macOS, Windows
- Core dependencies: typer ≥ 0.15.0, rich ≈ 13.0, pandas ≈ 2.0, requests ≈ 2.32, dendropy ≥ 4.6 and < 6.0 (both 4.6.x and 5.0.x verified with the full test suite; benchmarks in this paper used 4.6.4), scipy ≈ 1.11, numpy ≥ 1.24, pydantic ≈ 2.5, PyYAML ≥ 6.0

Install from source (to review the exact submission version, run git checkout v1.0.3 after cloning; the development install below is not the preferred reproduction path):

```
git clone https://github.com/ZengZichao/PyiTOL.git
cd PyiTOL
pip install -e ".[dev]"
```

#### S2.3 Docker deployment

PyiTOL provides a Docker image for isolated environments (HPC clusters or cloud platforms):

```
# Build image using the included Dockerfile (build from the archived v1.0.3 source for the reviewed version)
docker build -t pyitol:latest .

# Run container, mounting a local data directory
docker run -v $(pwd)/data:/data pyitol:latest \
  pyitol template create color-strip --tree /data/tree.nwk \
  --taxonomy /data/tax.csv --column Phylum -o /data/output.txt
```

#### S2.4 Long-term maintenance

PyiTOL is maintained by a team at Shanghai Jiao Tong University. Issues and feature requests are collected via GitHub Issues. Maintenance plans include adding support for new iTOL dataset types, performance optimization, documentation updates and real API integration tests. The authors commit to maintaining software availability for at least two years following publication.

### S3 | Algorithm Details

#### S3.1 Polytomy handling

iTOL itself supports Newick/Nexus trees containing polytomies, and PyiTOL's parser and API upload path pass such trees normally. For monophyly determination, when the LCA is a polytomy and members occupy only a subset of its child branches, the relationships among child branches are unresolved and no definitive monophyly/paraphyly/polyphyly call is justified. PyiTOL provides three user-selectable modes (strict and data-insufficient currently implement identical behaviour; relaxed is explicitly heuristic, see below):

- **strict mode** (default): Such unresolved cases return `data_insufficient`, with the `polytomy_details` field recording the polytomy position and the fraction of child branches occupied (e.g., `2/4`). This mode avoids the forced polyphyly over-calling of naive implementations on polytomies. If members span every child clade of the polytomy, the polytomy is the true LCA and the standard logic of main-text §2.3 applies.

- **relaxed mode (--polytomy-mode relaxed):** The same unresolved cases are classified as paraphyletic rather than data-insufficient — an explicitly heuristic review flag, not a resolved-topology determination, with polytomy_details likewise recorded. This mode suits taxonomic-revision scenarios, avoiding the loss of genera/families that still require revision merely because the branch structure is unresolved. Taxonomically, when a polytomy contains different taxonomic units (e.g., (A, B, C) belonging to different genera), classifying it as "paraphyletic" rather than "polyphyletic" is rigorous because an unresolved topology does not imply that the groups are evolutionarily unrelated. This conservative strategy aligns with the phylogenetic classification criteria of Wiley & Lieberman [21]: when topological relationships are insufficiently resolved, overly aggressive taxonomic decisions should be avoided. Note that this label is a pragmatic convention rather than a strict application of Farris's (1974) definitions [15] — an unresolved topology is, strictly speaking, neither paraphyletic nor polyphyletic but simply unresolved.
- **data-insufficient mode (--polytomy-mode data-insufficient):** Explicitly returns data_insufficient. Its behaviour is currently identical to strict mode — the two options are aliases in the current implementation. The separate name is retained for interface compatibility and as a semantic marker for downstream pipelines ("treat all unresolved topologies uniformly as data insufficient"); no behavioural difference should be assumed between them in this version.

#### S3.2 Pseudocode

```
Assumptions. T is interpreted as rooted as supplied and must carry unique tip labels (duplicate labels are rejected by the
validator before classification; §S8). S is the expected member set; the located members are P = S ∩ U (collected in step 1),
the missing members are M = S \ U, and when P = ∅ the topology is not assessed (unknown). All set operations below are over tip
identifiers.

Algorithm 1: check_monophyly_full(T, G, S, tip_to_group, polytomy_mode)
Input: Tree T, taxon name G, member set S, tip-to-group mapping tip_to_group
       (may be None), polytomy mode ∈ {strict, relaxed, data-insufficient}
Output: Classification status (monophyletic | paraphyletic | polyphyletic |
        incomplete_sampling | data_insufficient | unknown)
        and auxiliary information (lca_node, extra_nodes, missing_nodes,
        subgroups, polytomy_details)

 1. tip_nodes ← {find_tip_node(T, m) | m ∈ S ∧ find_tip_node(T, m) ≠ None}
 2. if |tip_nodes| = 0 then
 3.     return unknown(M = S)                          ▷ all members absent
 4. lca ← safe_mrca(tip_nodes)                        ▷ T rooted as supplied; flag only suppresses DendroPy re-rooting
 5. L ← {node.taxon.label | node ∈ leaf_iter(lca)}
 6. E ← L \ S      ▷ extra tips under the LCA
 7. M ← S \ L      ▷ expected members absent from the tree
 8. if is_polytomy(lca) and members_occupancy(lca, S) < total_children(lca) then
 9.     if M ≠ ∅ then return incomplete_sampling(M)
10.     if polytomy_mode ∈ {strict, data-insufficient} then
11.         return data_insufficient(E, polytomy_details)
12.     else                                            ▷ relaxed
13.         return paraphyletic(E, polytomy_details)
14. if E = ∅ and M = ∅ then
15.     return monophyletic(lca)
16. if M ≠ ∅ then
17.     return incomplete_sampling(M)        ▷ sampling absence takes precedence
18. if tip_to_group provides a usable mapping then
19.     if every node in E maps to G then
20.         status ← monophyletic              ▷ nested monophyly, extras = E
21.     else if some node in E maps to a different group then
22.         status ← mixed_children_classify(lca, S)   ▷ ≤1 mixed child → para; ≥2 → poly
23.     else
24.         status ← data_insufficient(E)     ▷ E contains unmapped nodes
25. else
26.     status ← mixed_children_classify(lca, S)       ▷ pure topology when unmapped
27. if status = polyphyletic then
28.     subgroups ← find_subgroups(tip_nodes)  ▷ maximal all-member clades, one post-order pass
29. return (status, lca, E, M, subgroups, polytomy_details)
```

Implementation notes. The algorithm is implemented in _check_monophyly_full() in core/monophyly.py. _find_tip_node() locates each member; a module-level _tip_index_cache caches the first-built {label: node} index to avoid repeated traversal (the --low-memory global option disables this cache, replacing indexed lookup with linear scanning). _safe_mrca() temporarily sets tree.is_rooted = True to unify the rooted state and avoid DendroPy's non-deterministic re-rooting of unrooted trees, restoring the original state in a finally block. PyiTOL does not infer or modify a biological root: every ancestor-based call is interpreted relative to the root as supplied by the user (main text §2.1); the temporary is_rooted flag only suppresses DendroPy's non-deterministic re-rooting side effect.

#### S3.3 Complexity analysis

Let n be the number of leaves and k the number of members of the taxon to be checked. For a single determination, LCA computation and descendant-set construction are O(n), and constructing the descendant-tip set L of the LCA together with the difference set E = L \ P costs O(|L|) ⊆ O(n), not O(k). The module-level tip-index cache reduces repeated member localization from O(n·k) to O(n + k) (O(n) to build the index once, then O(1) per lookup). The cache is protected by threading.Lock and cleared on each public entry call via clear_tip_index_cache(), ensuring thread safety and memory control. Subgroup decomposition is a single deterministic post-order traversal: leaf and member counts are accumulated bottom-up in O(n), after which the maximal member-pure clades are collected in a second O(n) pass, i.e., O(n) per polyphyletic group, independent of k. (An earlier greedy MRCA-expansion implementation with $O(k^2 \cdot n)$ cost per seed was replaced during this study's simulation-based accuracy evaluation after it was found to fragment genuine clades into singletons.) Across a batch of g groups the total cost is O(g·n) in the worst case; the shared tip-index keeps member localization at O(1) per member after a single O(n) index build, so the main text's per-group O(n) statements are not claims about the whole batch.

In real biological data, k is usually much smaller than n. Taking the GTDB R232 bac120 reference tree as an example, the median genus size among the 17,294 checked genera (those with two or more genomes in the derived taxonomy; singleton genera are trivially monophyletic) is 4 genomes, and the largest genus contains 2,311 genomes; at this typical scale the algorithm behaves approximately linearly. Note that the synthetic horizontal-comparison benchmarks deliberately use large groups with k = n/20 (horizontal comparison) and k = n/100 (extreme scale) to stress-test the subgroup-decomposition path; their k values are not representative of real taxonomic data.

#### S3.4 Subgroup decomposition

_find_subgroups() partitions a polyphyletic group into its maximal monophyletic subgroups. The key observation is that a subgroup is monophyletic if and only if its member set equals the complete leaf set of its LCA — equivalently, it is the leaf set of some tree node whose descendants are all members of the group. The maximal such clades are mutually disjoint, cover every member of the group, and constitute the unique minimum-cardinality partition of the group into monophyletic subgroups. PyiTOL collects them in a single deterministic post-order pass that accumulates per-node leaf counts and member counts and emits every member-pure clade whose parent is not member-pure; the procedure is therefore exact, order-independent, and O(n) per group. Proof of uniqueness and minimality: call a node member-pure when its descendant-tip set contains only group members. Maximal member-pure clades are pairwise disjoint, because nested member-pure clades collapse into the outer one; they cover every located member, because each member belongs to at least one member-pure clade (its own leaf or a member-pure ancestor) and hence to a maximal one. Any partition of the group into monophyletic subgroups has each part contained in some maximal member-pure clade, so no partition can use fewer parts, and the partition is unique. Boundary conditions: single-member subgroups are permitted; zero-length branches do not affect the leaf-set argument; polytomies are handled by the modes of §S3.1; duplicate tip labels are rejected upstream (§S8). For taxonomic-revision workflows the returned subgroups serve as candidate boundaries for re-describing or re-evaluating the affected taxa; confirming them as independent lineages (e.g., for taxon elevation) requires branch support, multi-tree agreement and external taxonomic evidence.

#### S3.5 Built-in special identifiers

PyiTOL ships four built-in special taxonomic identifiers — `LUCA`, `LACA`, `LBCA` and `ROOT` — which resolve to the last universal/common ancestor of all Bacteria and Archaea, all Archaea, all Bacteria, and the tree root (all leaves), respectively; they are provided as configurable defaults for domain-wide monophyly checks and are resolved at runtime via `_SPECIAL_IDENTIFIERS` in `core/monophyly.py`.

### S4 | Functional Workflow Test Checklist

The automated test suite comprises 72 test modules with 1,704 test cases (all passing in CI). The end-to-end functional checklist is:

| # | Workflow area | Test modules (representative) | Verified scenarios |
|---|---|---|---|
| 1 | CLI startup & banner | test_main_entry.py, cli/test_main.py, cli/test_apps.py | version banner, help text, subcommand routing, self-test |
| 2 | Configuration system | cli/test_config.py, test_config_merge* | multi-file YAML/JSON merge, CLI > config > default precedence, parameter aliases |
| 3 | Tree parsing | core/test_parser.py | Newick/Nexus auto-detection, BOM-prefixed files, multi-tree strategies (ask/first/last/split), large-tree warnings |
| 4 | Input validation | core/test_validator.py | hex/RGB/HSL/CSS-named colors, delimiter-conflict localization, numeric ranges, special characters in IDs, node-ID consistency |
| 5 | Taxonomy extraction | cli/test_taxonomy.py, cli/test_taxonomy_style.py | embedded-label and GTDB-format extraction, rank member lists, special identifiers (LUCA/LACA/LBCA/ROOT) |
| 6 | Monophyly classification | core/test_monophyly.py (66 tests) | all Table S2 scenarios, polytomy modes, nested monophyly, incomplete sampling, polyphyletic subgroup decomposition (including sibling-merge and anti-fragmentation regression tests added in this study) |
| 7 | Template generation | templates/test_*.py, cli/test_template_*.py | all 31 registered types, header/column validation, unified-entry ↔ subcommand consistency, LRU cache |
| 8 | API client | api/test_client.py, api/test_client_robustness*.py, api/test_error_translation.py | ZIP packaging, response parsing, retry/backoff, HTML error-page detection, offline mock integration, bilingual error translation |
| 9 | Session snapshot & replay | utils/test_session*.py, cli/ replay tests | snapshot schema (Pydantic), API-key redaction, --dry-run, step-by-step replay |
| 10 | Concurrency & interruption | test_concurrency.py, utils/test_shutdown*.py | thread-safe tip-index cache, SIGINT clean exit via subprocess signal tests |
| 11 | Exception hierarchy | test_exceptions.py | structured error codes, bilingual message rendering |
| 12 | Realistic data | test_realistic_data.py, tests/regression/ | GTDB-scale inputs, previously reported regressions frozen as tests |

### S5 | Reproduction Script for Extreme-Scale Monophyly Analysis

The 100,000- and 200,000-leaf synthetic-tree batch monophyly-check results are summarized in main-text Table 5. The reproduction script is benchmarks/benchmark_pyiTOL_extreme.py. Run the following command to regenerate the corresponding data locally:

```
python benchmarks/benchmark_pyiTOL_extreme.py --sizes 100000,200000 --repeats 3
```

The script generates deterministic fully balanced binary trees by recursive halving (not a stochastic process) and executes batch determination on 100 taxa via pyitol.core.monophyly.check_monophyly; taxa are assigned by (i // 40) % 100 (n/100 members per taxon, spread in contiguous blocks of 40 leaves), producing a worst-case, nearly-all-polyphyletic workload. Each size is run --repeats times (default 3); the script reports the median wall-clock time plus peak tracemalloc memory and saves all per-replicate values (the main text reports the median together with the full min–max range across the three replicates) to benchmarks/extreme_scale_results.json. Runtime depends on CPU, memory, and system load, so exact times may vary across workstations.

### S6 | Per-Module Code Coverage

Table S1 (in Section S11 below) shows statement coverage for the main PyiTOL modules measured on 2026-08-26 from a single full test run of the PyiTOL 1.0.3 codebase (1,704 tests: 1,703 passed, 1 skipped) with pytest-cov; overall statement coverage is 86.5%. Figure S1, Table S1 and the corresponding alt text are generated from the same coverage.json artifact of this run. Uncovered lines concentrate in extreme boundary conditions (e.g., multithreading lock paths and strict/relaxed logic switching on high-arity polytomies) that are exercised by manually constructed regression tests (§S4). Coverage quantifies the extent of automated testing; it is not a direct measure of scientific correctness, and the frequency with which uncovered paths occur in real biological data is not measured here.

### S7 | Example Datasets, GTDB Case-Study Reproduction and Deployment Guidance

#### S7.1 Example datasets

The project provides both real and synthetic example datasets. Real GTDB R232 reference trees are redistributed with the repository under `data/gtdb_r232/` for convenience; they are used only for the biological case study in the manuscript and are not included in the PyiTOL pip package. The original GTDB R232 data are publicly available at https://gtdb.ecogenomic.org/downloads; suffixed GTDB taxa (e.g., phylum `Bacillota_I`) are treated as distinct taxa throughout, following GTDB's rank-tree convention:

- `ar53_r232.tree`: archaeal 53-marker-gene reference tree, 10,122 genomes
- `bac120_r232.tree`: bacterial 120-marker-gene reference tree, 189,801 genomes

The following derived files can be regenerated using `scripts/extract_gtdb_taxonomy.py` and the commands described in §S7.2:

- `*_taxonomy.csv`: six-rank taxonomy metadata inferred from internal-node labels
- `*_phylum_strip.txt`: PyiTOL-generated phylum-level color-strip templates
- `*_monophyly_genus.csv`: genus-level monophyly validation results

The real phylogenomic tree used in main-text §4.5 is shipped with the repository:

- `benchmarks/data/FigTree_withLACA_CLK_95CI.tree.recover`: relaxed molecular-clock tree of 700 archaeal and bacterial genomes, derived from the published dataset (see provenance below) — renamed and tip-label re-annotated by the present authors to embed six-rank taxonomy, topology unmodified (NEXUS; BEAST-style 95% CI annotations are ignored by the parser)
- Derived files regenerable via `scripts/extract_figtree_taxonomy.py` (`laca_tree_taxonomy.csv`) and `pyitol taxonomy monophyly --rank {genus|family|order|class|phylum|domain}` (laca_monophyly_<rank>_strict.csv)
- Cross-validation script against ete3/ete4: `scripts/cross_validate_laca.py`

**Data provenance of the 700-genome tree.** The tree was not inferred by the present authors: it was downloaded from the publicly archived supplementary data of Moody et al. [19] (Moody ERR, Williams TA, Álvarez-Carretero S, Szöllősi GJ, Pisani D, Lenton TM, Donoghue PCJ. 2025 The emergence of metabolisms through Earth history and implications for biospheric evolution. Phil. Trans. R. Soc. B 380: 20240097, DOI: 10.1098/rstb.2024.0097; supplementary data archived at figshare, dataset DOI: 10.6084/m9.figshare.27968166 [main-text reference 20]), in which a time-calibrated tree of life spanning Bacteria and Archaea was inferred under a relaxed molecular clock. It was deliberately chosen for this evaluation because its taxonomy is not curated to be monophyly-consistent, so non-monophyly signals are expected a priori. The dataset is licensed under CC BY 4.0 and is redistributed in accordance with that license, with attribution to [19,20] and with modifications indicated as required: the file was renamed and its tip labels were re-annotated by the present authors to embed six-rank taxonomic lineage, while the tree itself (topology, branch lengths, and node annotations) is unmodified; the unmodified original is retrievable from the figshare dataset [20] for comparison. All biological interpretation of the tree itself is due to [19], and the present work uses it solely as a public real-data case study for PyiTOL's classification engine; because the tip taxonomy was re-annotated by the present authors, the resulting calls are tree-relative candidates rather than an independent ground truth.

Synthetic examples are located in `examples/data/`:

- `synthetic_32tips.nwk` and `synthetic_32tips_taxonomy.csv`: small synthetic phylogenetic tree (32 tips) with three-rank taxonomy metadata, suitable for quick verification
- `synthetic_32tips_enriched.csv`, `synthetic_32tips_auto_taxonomy.csv`: extended metadata and an auto-extracted taxonomy example for the 32-tip tree
- `synthetic_130tips.nwk` and `synthetic_130tips_taxonomy.csv`: phylogenetic tree of Enterobacterales genomes with taxonomy metadata
- `synthetic_130tips_enriched.csv`: enriched metadata including GC content and genome size
- `synthetic_130tips_auto_taxonomy.csv`: auto-extracted taxonomy example
- `synthetic_130tips_amr.csv`: antibiotic-resistance gene presence/absence data
- `synthetic_4096tips.nwk` and `synthetic_4096tips_taxonomy.csv`: bacterial-genome phylogenetic tree simulated at GTDB scale with six-rank taxonomy metadata
- `synthetic_4096tips_traits.csv`: functional-gene presence/absence matrix

- `synthetic_5000tips.nwk`, `synthetic_5000tips_taxonomy.csv`, `synthetic_5000tips_traits.csv`: 5,000-tip synthetic tree with accompanying taxonomy metadata and trait matrix

Accuracy-benchmark artifacts (`benchmarks/accuracy_results/accuracy_summary.json`, per-replicate trees, taxonomies, and result CSVs for both the balanced and random regimes) and API-acceptance artifacts (`benchmarks/itol_acceptance/acceptance_results.json`, generated template files) are likewise included in the repository.

### S7.2 GTDB case-study reproduction commands

Reproduction of the GTDB-derived artifacts used in main-text §4.5 (taxonomy extraction, validation, color-strip templates and genus-level monophyly CSVs):

```
# 1. Extract terminal taxonomy information from GTDB reference tree internal-node labels
python scripts/extract_gtdb_taxonomy.py --tree data/gtdb_r232/ar53_r232.tree \
  --output data/gtdb_r232/ar53_r232_taxonomy.csv --rank genus \
  --taxa-output data/gtdb_r232/ar53_r232_genera.txt
python scripts/extract_gtdb_taxonomy.py --tree data/gtdb_r232/bac120_r232.tree \
  --output data/gtdb_r232/bac120_r232_taxonomy.csv --rank genus \
  --taxa-output data/gtdb_r232/bac120_r232_genera.txt

# 2. Validation and template generation (phylum-level color strip example)
pyitol validate --tree data/gtdb_r232/ar53_r232.tree \
  --taxonomy data/gtdb_r232/ar53_r232_taxonomy.csv
pyitol template create color-strip --tree data/gtdb_r232/ar53_r232.tree \
  --taxonomy data/gtdb_r232/ar53_r232_taxonomy.csv --column phylum \
  -o data/gtdb_r232/ar53_r232_phylum_strip.txt
pyitol template create color-strip --tree data/gtdb_r232/bac120_r232.tree \
  --taxonomy data/gtdb_r232/bac120_r232_taxonomy.csv --column phylum \
  -o data/gtdb_r232/bac120_r232_phylum_strip.txt

# 3. Genus-level monophyly validation (conservative polytomy mode)
pyitol taxonomy monophyly --tree data/gtdb_r232/ar53_r232.tree \
  --taxonomy data/gtdb_r232/ar53_r232_taxonomy.csv --rank genus \
  --taxa-file data/gtdb_r232/ar53_r232_genera.txt \
  --polytomy-mode relaxed -o data/gtdb_r232/ar53_r232_monophyly_genus.csv
pyitol taxonomy monophyly --tree data/gtdb_r232/bac120_r232.tree \
  --taxonomy data/gtdb_r232/bac120_r232_taxonomy.csv --rank genus \
  --taxa-file data/gtdb_r232/bac120_r232_genera.txt \
  --polytomy-mode relaxed -o data/gtdb_r232/bac120_r232_monophyly_genus.csv
```

Timings reproduced on a MacBook Pro (Apple M5): phylum-level color-strip generation about 0.7 s (ar53) and 8 s (bac120); genus-level monophyly validation 0.95 s (ar53, peak memory about 154 MiB) and about 17 s (bac120, 12–17 s across cold/warm disk cache; peak memory about 4.9 GiB). The commands above write into data/gtdb_r232/; when reproducing against the archived reference results, redirect outputs to a separate directory (e.g., repro/gtdb_r232/) so that the archived copies are not overwritten.

**Figure 5 colour mappings.** In main-text Figure 5, each branch colour is determined by the first letter of the phylum name through a fixed per-letter colourblind-safe palette; phyla sharing the same initial letter are coloured identically, and the letter-to-colour mapping is identical in both panels (tips whose phylum cannot be parsed fall back to uniform grey #B0B0B0). Tip labels use a uniform dark colour (#333333), with taxonomic information carried solely by the colour strip. MRCA nodes of collapsed genera/orders are marked with yellow triangle symbols (#d4b86a); completeness is shown as star-shaped bubbles (CheckM teal #00A087, CheckM2 blue #3C5488; larger bubble = higher completeness, %), contamination as circle bubbles (CheckM red #B2182B, CheckM2 orange-red #D6604D; larger bubble = higher contamination, %), followed by a GC-content bar track (blue #4393C3, %) and a genome-size bar track (orange #E66101, log10 bp), with the outermost phylum colour strip matching branch colours. LEGEND blocks for the numeric layers are written into the corresponding iTOL template files; full interactive legends are available in the iTOL interface. Provenance of the quality indicators: completeness and contamination (CheckM and CheckM2), GC content and genome size are taken from the GTDB R232 metadata table redistributed with the repository; values shown for collapsed clades are arithmetic means over member genomes, with genomes missing a given metric excluded from that mean; the aggregation and per-layer inputs are implemented in the figure-generation script (A-正文图片/图 5_GTDB 物种树折叠.py).

### S7.3 Practical considerations for real-world deployment

PyiTOL sits downstream of phylogenetic analysis pipelines, where upstream tools such as RAxML [22], IQ-TREE [23], FastTree [24] or PhyloPhlAn [25] typically generate Newick trees, and GTDB-Tk [27,28], CheckM [26] or custom scripts generate taxonomic annotations (16S rRNA-based studies may equally use taxonomy databases such as SILVA [29], integrated through the same column structure). To ensure reliable PyiTOL monophyly determinations, we recommend the following preprocessing at the upstream stage:

1. **Remove or collapse low-support branches.** Branches with very low bootstrap support (e.g., < 50) often represent unresolved topological noise. First collapse them into polytomies using RAxML/IQ-TREE -collapse options or Newick tools such as nw_topology

or phylopy, then run PyiTOL with --polytomy-mode relaxed as a heuristic review aid (relaxed is a pragmatic convention rather than a resolved-topology determination, §S3.1), reducing the risk that noise is read as polyphyly.

2. **Unify terminal labels and metadata IDs.** MAG terminal labels often differ from IDs in taxonomy tables (e.g., containing spaces, version suffixes `.1`, or sample prefixes). We recommend using `pyitol validate` to pre-check node-ID consistency and cleaning labels via `taxonomy extract` or custom scripts.
3. **Handle polytomies and uncultured taxa.** For trees composed mainly of MAGs, many uncultured genera (e.g., `g__UBA123`, `f__JABCX01`) may appear in unresolved positions due to incomplete marker genes. `--polytomy-mode relaxed` reduces over-splitting, but results should still be combined with CheckM completeness/contamination or GTDB quality scores to filter low-quality genomes.
4. **Partition ultra-large trees.** When tree size exceeds 100,000 leaves, we recommend cutting the tree into subtrees by phylum/class and running monophyly checks separately, or generating templates only for target taxa rather than processing the whole tree at once, to reduce memory peaks and improve iteration speed. The ~100,000-leaf threshold is an empirical heuristic, not a universal rule; note also that partitioning can hide non-monophyly spanning partition boundaries — where feasible, re-check boundary taxa on a larger subtree or the unpartitioned tree.

These preprocessing recommendations also apply to downstream visualization: cleaned taxonomy tables and collapsed tree files can be uploaded directly to iTOL, avoiding repeated adjustments in the iTOL web interface.

### S8 | Input Validation and Robustness

#### S8.1 Implementation details

**BOM-safe file reading.** Tree and metadata files saved by some editors (including Windows Excel and Notepad) may carry an invisible UTF-8 Byte Order Mark (BOM, U+FEFF). The BOM is harmless for general text display but breaks Newick/Nexus parsers and CSV readers that expect files to start with specific characters, producing cryptic errors such as "illegal character" or "header mismatch". safe_read_text() in utils/io.py detects and removes a leading U+FEFF and decodes with a utf-8-sig fallback; the verified scope is UTF-8 (with or without BOM) — UTF-16 and other encodings are out of scope. The parser and validator first read file contents as strings and then call DendroPy's get_from_string(), avoiding parse failures caused by BOM-prefixed files.

**Multi-tree file handling.** load_tree() supports four named multi-tree strategies — ask, first, last, and split (split and all are synonyms) — plus explicit --tree-index selection; the project maintains deterministic behaviour and provides no random strategy. When the strategy is ask and the environment is non-interactive, it automatically falls back to first and prints a warning, preventing unexpected failures in scripts or CI. Users can explicitly specify the strategy with --multi-tree-mode first --quiet to suppress warning messages.

**Color, delimiter, and range validation.** `validate_color_code()` supports hexadecimal, RGB/RGBA, HSL/HSLA, and 113 CSS named colors, returning an `(is_valid, suggestion)` tuple. `check_delimiter_conflict()` uses vectorized string operations to locate cells that conflict with the chosen delimiter. `check_numeric_range()` includes biologically reasonable ranges for GC content, bootstrap support values, branch lengths, etc.

**Signal-safe interruption.** The `GracefulShutdownContext` context manager in `utils/shutdown.py` wraps SIGINT/SIGTERM signal handling and provides `check_shutdown()`, `update_progress()`, `register_temp()`, and `register_handle()` interfaces. Module-level function `register_signal_handlers()` registers signal handlers at CLI entry startup; `register_temp_file()` and `register_file_handle()` provide global resource registration. When the user presses Ctrl+C, a clean exit is triggered; test scripts verify this mechanism by sending SIGINT via Python `subprocess`, eliminating the system dependency on GNU `timeout`.

#### S8.2 Robustness test results

The robustness mechanisms above were exercised by dedicated test cases covering the following anomalous-input scenarios, as summarised below; each scenario is covered by regression tests (1,704 tests in the full suite; relevant cases in tests/core/test_parser.py, tests/core/test_validator.py and tests/utils/):

- **BOM files:** UTF-8-BOM encoded tree files parse correctly (UTF-16 is out of scope; §S8.1).
- **Empty and non-existent files**: `TreeParser` raises `TreeParseError` for empty files and logs a warning returning an empty parser for non-existent files.
- **Missing members**: Members not present in the tree are faithfully recorded as `missing_nodes`; groups containing absent members are flagged as `incomplete_sampling`, with a summary warning emitted instead of silently dropping them.
- **Special characters**: `check_special_characters_in_ids()` detects spaces, tabs, commas, semicolons, and vertical bars in node IDs.

- **Multi-tree files**: The `ask` strategy automatically falls back to `first` in non-interactive environments.
- **Invalid colors and delimiter conflicts**: Return structured error codes and remediation suggestions.
- **API errors**: iTOL errors such as `invalid project name` are automatically translated into Chinese operation suggestions, and empty server responses produce an explicit diagnostic marker rather than a blank error.

### S9 | Bilingual Error Message Examples

The following examples show representative bilingual message formats produced by the current English/Chinese message catalog of PyiTOL 1.0.3 (message wording may evolve between versions; the blocks below are illustrative renderings rather than verbatim terminal captures):

**Example 1: Invalid iTOL project name**

```
# Original iTOL API response (English):
ERROR: invalid project name

# PyiTOL translated output (PYITOL_LANG=zh):
[ERROR] iTOL API 错误: invalid project name
[提示] 项目名称无效。请登录 iTOL 网站确认项目名是否已创建，并检查拼写是否正确。
```

**Example 2: Tree parse error with BOM**

```
# Without PyiTOL (raw DendroPy error):
dendropy.dataio.newickreader.NewickReader: Unexpected token: '\ufeff('

# PyiTOL bilingual output:
[ERROR] 树文件解析失败 / Tree file parse error
[详情] 检测到 BOM 标记，已自动移除并重新解析。
[Details] BOM marker detected and automatically removed; retrying parse.
```

**Example 3: Color validation suggestion**

```
# Input: color code "#GG0000"
# PyiTOL output:
[WARNING] 无效颜色代码 / Invalid color code: '#GG0000'
[建议] 您是否想使用 '#FF0000' (red)?
[Suggestion] Did you mean '#FF0000' (red)?
```

### S10 | Declaration on the Use of AI-Assisted Tools

In accordance with the journal's policy on the acceptable use of large language models (LLMs), the authors declare that an LLM-based assistant was used for two purposes during the preparation of this work: (i) language editing and polishing of the manuscript text, and (ii) assisting in the writing of the source code (including implementation of algorithms and helper functions). All scientific content was conceived, executed, and verified by the authors. The authors reviewed and revised all AI-generated suggestions (both code and text) and take full responsibility for the correctness and integrity of the code and the final manuscript.

### S11 | Supplementary Tables

**Table S1 | Per-module statement coverage**

| Module | Coverage | Notes |
|---|---|---|
| core | 75.1% | Boundary conditions in validator.py/taxonomy.py and high-arity polytomy paths; exercised by regression tests (§S4) |
| api | 92.2% | Retry/polling and export-validation paths covered; live-server calls mocked in CI |
| templates | 90.2% | Uncovered lines concentrated in rarely used schema branches |
| utils | 95.7% | IO/session/color utilities near-complete |
| cli | 88.5% | Command wiring and option parsing |
| Overall | 86.5% | Measured 2026-08-26 on the PyiTOL 1.0.3 codebase (1,704 tests: 1,703 passed, 1 skipped); --cov-fail-under=80 |

**Table S2 | Monophyly classification sandbox test scenarios and results**

Eleven boundary-condition scenarios covering binary trees, polytomies, single-member groups, whole-tree groups, nested monophyly, paraphyly, polyphyly, missing members, data insufficiency and unknown status. All scenarios passed; classification results matched the formalization of main-text §2.3. Results are reproducible via `tests/core/test_monophyly.py`.

| Scenario | Tree topology | Member set | Mode | Expected | Observed | Result |
|---|---|---|---|---|---|---|
| Binary-tree monophyly | ((A,B),(C,D)) | {A,B} | strict | monophyletic | monophyletic | ✓ |
| Classic paraphyly (nested extra) | ((A,(B,X)),C) | {A,B} | strict | paraphyletic | paraphyletic | ✓ |
| Unresolved polytomy (strict) | (A,B,(C,D)) | {A,B} | strict | data_insufficient | data_insufficient | ✓ |
| Conservative polytomy call (relaxed) | (A,B,(C,D)) | {A,B} | relaxed | paraphyletic | paraphyletic | ✓ |
| Single-member group | ((A,B),(C,D)) | {A} | strict | monophyletic | monophyletic | ✓ |
| Polyphyly detection | ((A,C),(B,D)) | {A,B} | strict | polyphyletic | polyphyletic | ✓ |
| Whole-tree group | ((A,B),(C,D)) | {A,B,C,D} | strict | monophyletic | monophyletic | ✓ |
| Nested monophyly | (((A,(C,D)),B),E) | {A,B} (C, D also in G) | strict | monophyletic | monophyletic | ✓ |
| Unmapped extra node | (((A,(C,D)),B),E), D unmapped | {A,B} | strict | data_insufficient | data_insufficient | ✓ |
| Empty member list | (A,B) | {} | strict | unknown | unknown | ✓ |
| Missing member (incomplete sampling) | (A,B) | {A,X} | strict | incomplete_sampling | incomplete_sampling | ✓ |

Note: for the 4-furcating tree `(A,B,C,D);` with G1={A,B}, G2={C}, G3={D}, strict and data-insufficient modes return `data_insufficient` (no forced classification of unresolved topology), whereas relaxed mode conservatively classifies the group as paraphyletic and records that members occupy 2/4 child branches in `polytomy_details`. In `((A,(B,X)),C)` the member set {A,B} has a single mixed child of its LCA with extra tip X nested inside it (at-most-one-mixed-child criterion). In `(A,B,(C,D))` the root is trifurcating, so the LCA is a polytomy whose members occupy only 2 of 3 child branches; strict mode therefore returns `data_insufficient`. In the polyphyly example `((A,C),(B,D))`, both children of the LCA are mixed children, satisfying the ≥2 mixed-children criterion.

**Table S3 | iTOL live-server acceptance of PyiTOL-generated templates (details)**

Each of the 31 types was generated by PyiTOL's `TemplateGenerator` against the 32-tip example tree and uploaded to the live iTOL server via PyiTOL's API client (batch uploader, ZIP packaging; iTOL v7.6, tested 2026-08-04); acceptance = server returned `SUCCESS` with a tree ID. Final result: 22/31 accepted (71%). Reproducible via `benchmarks/benchmark_itol_acceptance.py` (raw responses: `benchmarks/itol_acceptance/acceptance_results.json`).

| Status | Count | Types | Server response |
|---|---|---|---|
| Accepted | 22 | alignment, binary, boxplot, branch/tree_colors, collapse, color_strip, connection, domains, externalshape, gradient, heatmap, labels, linechart, multi_bar, pie, popup_info, range, simple_bar, spacing, style, symbol, text | SUCCESS with numeric tree ID |
| Rejected — explicit batch-mode limitation | 1 | image | "WARNING: DATASET_IMAGE is not supported in batch mode" |
| Rejected: not identified by the batch uploader — failure causes (i)–(iv) in main-text §4.4 | 8 | arrows, manual, meme, placement, prune, tanglegram, timescale, treestyle | "ERR 5 … Failed to identify the type of file" |

Interpretation of the nine rejections (all server-side, not formatting defects): (1) DATASET_IMAGE is documented as unsupported by the batch uploader; (2) DATASET_MANUAL has no template file by design (interactive drawing tools only, per iTOL help); (3) DATASET_PLACEMENT data are extracted from `.jplace` files rather than tabular datasets; (4) iTOL v7 DATASET_TANGLEGRAM requires the second tree embedded between `TANGLEGRAM_TREE`/`END_TANGLEGRAM_TREE` fields inside the dataset, a v6→v7 structural change not covered by the tabular id1/id2 mapping; (5) PRUNE, DATASET_TIMESCALE, DATASET_TREESTYLE, DATASET_ARROWS, and DATASET_MEME are applied through the web interface or control panel and are not identified by the batch uploader. All nine types remain fully generable by PyiTOL for manual upload through the iTOL web interface.

Testing notes. Two server-side behaviours required test-data adaptation and are worth reporting for other API users: (a) the batch uploader converts underscores in unquoted Newick labels to spaces, so annotation dataset IDs must use the space form (or the tree labels must be quoted); (b) the COLLAPSE dataset, in contrast, matches IDs against the raw Newick tokens — with quoted labels (underscores preserved) COLLAPSE fails with ERR 8 while annotation datasets succeed, and with unquoted labels the opposite holds. We therefore used unquoted Newick plus space-form IDs for annotation types and underscore (raw-token) IDs for COLLAPSE. Test templates additionally had to avoid non-existent parameter names (e.g., `BAR_ALIGN`, `INNER_RADIUS`, `BOX_COLOR` are rejected as unknown variables by the current server).

**Table S4 | Simulation rule conformance: merged confusion matrix (predicted rows × engineered columns)**

From 16 replicates — balanced regime (2,048 and 8,192 leaves × 5 seeds, 3,800 groups) and random-bifurcating regime (2,048 and 4,096 leaves × 3 seeds, 589 groups) — 4,389 groups were scored; all off-diagonal entries are zero. The engineered paraphyletic column merges the shallow construction (one whole subclade removed) and the deeply nested variant (extras nested two levels below the LCA), whose predicted label is identical. Reproducible via `benchmarks/benchmark_accuracy_simulation.py` (raw data: `benchmarks/accuracy_results/accuracy_summary.json`).

| Predicted Engineered | monophyletic | paraphyletic | polyphyletic | incomplete_sampling | data_insufficient |
|---|---|---|---|---|---|
| monophyletic | 2,401 | 0 | 0 | 0 | 0 |
| paraphyletic | 0 | 797 | 0 | 0 | 0 |
| polyphyletic | 0 | 0 | 397 | 0 | 0 |
| incomplete_sampling | 0 | 0 | 0 | 397 | 0 |
| data_insufficient | 0 | 0 | 0 | 0 | 397 |

**Table S5 | Cross-validation summary on the 700-genome phylogenomic tree (genus level)**

| Comparison | Result |
|---|---|
| PyiTOL vs ete4, binary (mono / non-mono) agreement | 409/409 (100%) |
| PyiTOL vs ete3, binary agreement on multi-member groups (≥2 members) | 23/23 (100%) |
| PyiTOL vs ete3, all 409 groups | 23/409 — the 386 discrepancies are all single-member genera that ete3 labels polyphyletic by design |
| Three-way discordance | *Thermococcus*: PyiTOL = paraphyletic, ete4 = paraphyletic, ete3 = polyphyletic (nested single extra tip *Palaeococcus*; at-most-one-mixed-child case) |
| Non-monophyletic genera detected | *Sulfolobus* (polyphyletic, decomposed into 2 subgroups), *Archaeoglobus* (paraphyletic, 2 nested extra tips), *Thermococcus* (paraphyletic, 1 nested extra tip) |

**Table S6 | Matched-scope full-mode benchmark (pre-parsed in-memory tree)**

To address the unequal timing scopes of main-text Table 3 (PyiTOL's end-to-end path includes disk re-parsing while DendroPy reuses a pre-parsed tree), PyiTOL's full 10-field mode was additionally timed on a pre-parsed in-memory tree with taxonomy loading excluded from the timed loop (`pyitol_inmem` mode of `benchmarks/benchmark_monophyly_comparison.py`); the tip-index cache is rebuilt inside each timed run, so index construction is still charged to the timed path. The Boolean-only lightweight variant (`pyitol_light`) shares the DendroPy control's algorithmic path (same `mrca()` + descendant-set comparison, same `taxon_namespace.get_taxon()` label resolution, `is_rooted = True`) and is included as the equal-task reference; its ~17% advantage over the DendroPy column at 50,000 leaves reflects library-internal overhead rather than algorithmic differences, both runs being dominated by the per-query linear label scan. Medians of 3 replicates; same trees and taxonomy as main-text Table 3.

| Leaves | PyiTOL full (end-to-end, Table 3) | PyiTOL full (matched scope, no parsing) | Speed-up | PyiTOL light (Boolean-only) | DendroPy control |
|---|---|---|---|---|---|
| 1,000 | 409.6 ms | 350.7 ms | 1.17× | 58.3 ms | 61.8 ms |
| 10,000 | 4.74 s | 3.73 s | 1.27× | 2.33 s | 2.33 s |
| 50,000 | 23.47 s | 18.97 s | 1.24× | 53.06 s | 63.99 s |

**Table S7 | PyiTOL template generation scalability benchmark**

| Leaves | Color strip (time) | Color strip (memory) | Heatmap (time) | Heatmap (memory) | Simple bar (time) | Simple bar (memory) |
|---|---|---|---|---|---|---|
| 100 | 3.7 ms | 230.0 KiB | 3.7 ms | 23.5 KiB | 3.5 ms | 19.3 KiB |
| 1,000 | 36.4 ms | 1.9 MiB | 38.3 ms | 293.0 KiB | 35.9 ms | 269.4 KiB |
| 10,000 | 357.9 ms | 18.3 MiB | 385.4 ms | 2.9 MiB | 388.3 ms | 2.7 MiB |
| 50,000 | 2.01 s | 92.0 MiB | 2.07 s | 14.7 MiB | 1.90 s | 13.6 MiB |

Note: Values are medians of 10 replicates per size (per-replicate min–max ranges are available in `benchmarks/template_scalability_results.json`). All three template types are measured under an identical methodology: a warm-up pass primes lazy imports and first-call caches before any measurement, and every reported memory value is a tracemalloc incremental allocation since `tracemalloc.start()` (no interpreter baseline is included in any cell). The color-strip template maintains a color map for each category, which explains its higher memory footprint; occasional slow outliers from OS scheduling make medians more representative than means.

**Table S8 | Comparison of output information content for monophyly determination**

| Tool | Returned fields | Main output content |
|---|---|---|
| PyiTOL | 10 | Classification status, LCA node, extra-node set, missing-node set, polyphyletic subgroups, polytomy details |
| ete3 | 3 | is_monophyletic (Boolean), clade type (monophyletic/paraphyletic/polyphyletic), set of extra leaves breaking monophyly |
| ete4 | 3 | Same as ete3 (ETE v4 rewrite; check_monophyly() returns the same 3-tuple) |
| DendroPy | 1 | MRCA node reference (monophyly requires manual descendant-set comparison) |

**Table S9 | Complete inventory of the 31 registered iTOL template types**

Canonical type names (with accepted aliases), iTOL header identifiers, and required data columns, as registered by the @register_template plugin system (no peer-reviewed iTOL v7 publication exists yet; batch acceptance was empirically validated for 22 of the 31 registered types by live-server uploads — Table S3 and main-text §4.4 give the per-type outcomes and failure-cause categories, and the remaining registered types are experimental pending the upgrades listed in main-text §5.2). The "Canonical type" column shows the user-facing CLI type names; the registry-internal canonical keys additionally carry a dataset_ prefix for annotation types (e.g., CLI name color_strip ↔ registry key dataset_colorstrip; multi_bar ↔ dataset_multibar; pie ↔ dataset_piechart; symbol ↔ dataset_symbols; connection ↔ dataset_connections; branch/tree_colors ↔ dataset_tree_colors). Both forms resolve to the same schema via the alias map.

| # | Canonical type | iTOL header | Required columns |
|---|---|---|---|
| 1 | alignment | DATASET_ALIGNMENT | id, alignment |
| 2 | arrows | DATASET_ARROWS | id, position |
| 3 | binary | DATASET_BINARY | id |
| 4 | boxplot | DATASET_BOXPLOT | id, minimum, q1, median, q3, maximum |
| 5 | branch / tree_colors | TREE_COLORS | id, type, color |
| 6 | collapse | COLLAPSE | id |
| 7 | color_strip | DATASET_COLORSTRIP | id, color, value |
| 8 | connection | DATASET_CONNECTION | id1, id2 |
| 9 | domains | DATASET_DOMAINS | id, from, to, type |
| 10 | externalshape | DATASET_EXTERNALSHAPE | id |
| 11 | gradient | DATASET_GRADIENT | id, value |
| 12 | heatmap | DATASET_HEATMAP | id |
| 13 | image | DATASET_IMAGE | id, image_file |
| 14 | labels | LABELS | id, label |
| 15 | linechart | DATASET_LINECHART | id, position, value |
| 16 | manual | DATASET_MANUAL | — |
| 17 | meme | DATASET_MEME | id, start, end, name |
| 18 | multi_bar | DATASET_MULTIBAR | id |
| 19 | pie | DATASET_PIECHART | id |
| 20 | placement | DATASET_PLACEMENT | id, position, count |
| 21 | popup_info | POPUP_INFO | id, title, content |
| 22 | prune | PRUNE | id |
| 23 | range | DATASET_RANGE | id, label, color |
| 24 | simple_bar | DATASET_SIMPLEBAR | id, bar_height |
| 25 | spacing | SPACING | id, factor |
| 26 | style | DATASET_STYLE | id, type, color |
| 27 | symbol | DATASET_SYMBOL | id, type, value, color |
| 28 | tanglegram | DATASET_TANGLEGRAM | id1, id2 |
| 29 | text | DATASET_TEXT | id, text |
| 30 | timescale | DATASET_TIMESCALE | time_point, label |
| 31 | treestyle | DATASET_TREESTYLE | — |

Types 1–27 are dataset types; types 6, 22, 25 (COLLAPSE, PRUNE, SPACING) plus treestyle form the four tree-structure types (TREE_COLORS is counted among the 27 dataset types because it is uploaded as an annotation dataset). The highlight feature is implemented by reusing the `style` type.

### S12 | Supplementary Figures

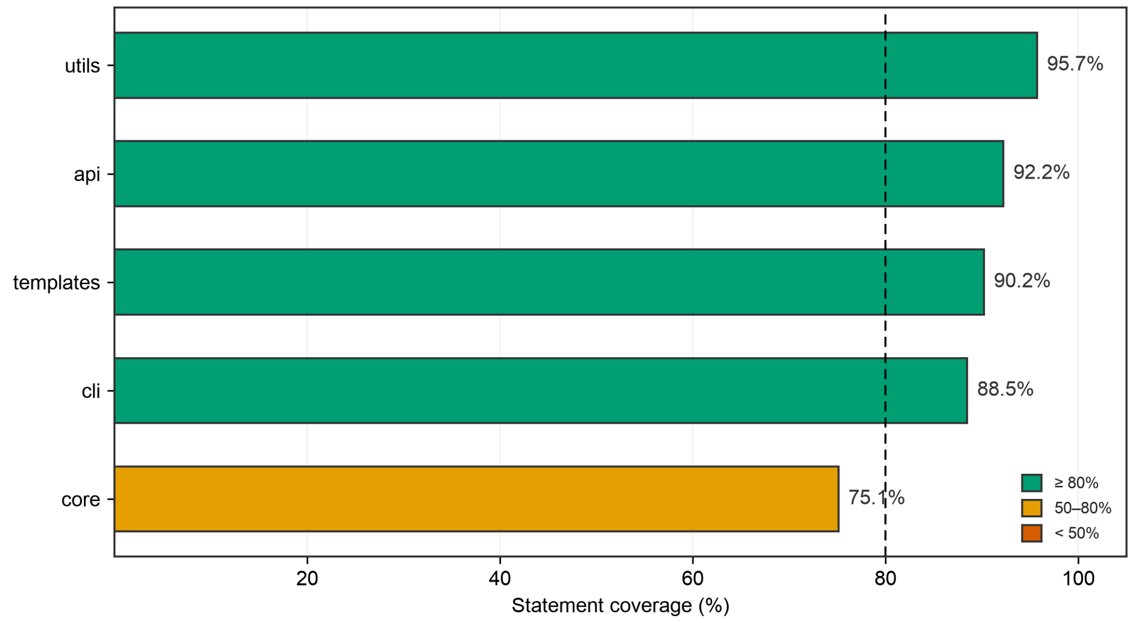


**Figure S1 | Test-coverage module distribution.** Horizontal bar chart showing automated-test statement coverage of the main modules, sorted in descending order of coverage (auto-generated from the coverage.json of the 2026-08-26 full test run; values identical to Table S1). Color coding indicates coverage sufficiency: green for modules with coverage ≥ 80%, yellow for 50%–80%, and red for < 50%. Overall statement coverage is 86.5%. Lower coverage in the core module is concentrated in extremely complex boundary conditions in validator.py and taxonomy.py — such as strict/relaxed logic switching on large numbers of child branches and exception-handling paths in multithreaded locks (all 113 defined CSS named colors are fully covered by parametrized tests). These scenarios are very rare in real biological data and have been verified by manually constructing boundary cases that are frozen as regression tests. Detailed module coverage data are provided by the CI --cov-report=xml artifact.

Alt text: Horizontal bar chart of automated-test statement coverage per main module (utils 95.7%, api 92.2%, templates 90.2%, cli 88.5%, core 75.1%), colour-coded by coverage sufficiency (green >=80%, yellow 50-80%, red <50%) and ranked by descending overall coverage; dashed line marks the 80% CI coverage threshold.

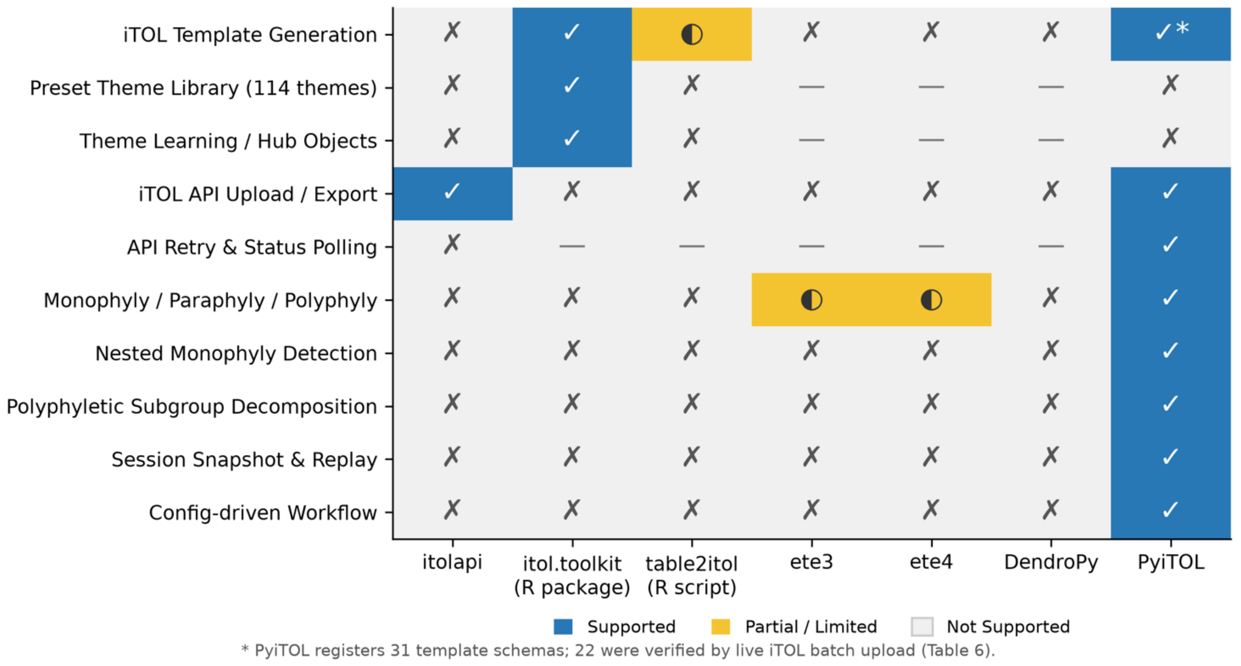


**Figure S2 | Feature-coverage comparison of PyiTOL with existing iTOL companion tools.** Heat-map style feature matrix comparing ten capabilities (including itol.toolkit's preset theme library and theme learning) across itolapi, itol.toolkit (R package), table2itol (R script), ete3, ete4, DendroPy, and PyiTOL. Blue = supported (✓), yellow = partial/limited (◐), gray = unsupported (✗) or not applicable (—, used only in the "API retry & status polling" row for tools without an API client). The PyiTOL column is highlighted with a dark border, showing its complete coverage of three-way monophyly classification, nested detection, polyphyletic subgroup decomposition, session snapshots, and config-driven workflows.

Alt text: Color-coded feature-coverage comparison matrix (itolapi, itol.toolkit R, table2itol R, ete3, ete4, DendroPy vs PyiTOL) across eight capabilities, with check, half-circle, cross and dash symbols ensuring grayscale/color-blind readability; no column highlighted; PyiTOL's template-generation cell carries an asterisk footnote (31 registered schemas, 22 verified by live batch upload).

**Figure S2 accessibility**: In the color-coded comparison matrix, each cell is annotated with a symbol (✓ for full support, ◐ for partial support, ✗ for not supported, — for not applicable) to ensure readability in grayscale printing and for color-blind readers.
